# From Typical Sequences to Typical Genotypes

**DOI:** 10.1101/079491

**Authors:** Omri Tal, Tat Dat Tran, Jacobus Portegies

**Affiliations:** Max-Planck-Institute for Mathematics in the Sciences, Leipzig, Germany Inselstrasse 22, D-04103 Leipzig

**Keywords:** typical sequences, typical genotypes, population entropy rate, population cross entropy rate, classification

## Abstract

We demonstrate an application of a core notion of information theory, that of typical sequences and their related properties, to analysis of population genetic data. Based on the asymptotic equipartition property (AEP) for non-stationary discrete-time sources producing independent symbols, we introduce the concepts of *typical genotypes* and *population entropy rate* and *cross entropy rate.* We analyze three perspectives on typical genotypes: a set perspective on the interplay of typical sets of genotypes from two populations, a geometric perspective on their structure in high dimensional space, and a statistical learning perspective on the prospects of constructing typical-set based classifiers. In particular, we show that such classifiers have a surprising resilience to noise originating from small population samples, and highlight the potential for further links between inference and communication.

## 1 Introduction

> We are drowning in information and starving for knowledge.
>
> — - John Naisbitt.

In this paper we identify several intrinsic properties of long stretches of genetic sequences from multiple populations that justify an information theoretic approach in their analysis. Our central observation is that long genotypes consisting of polymorphisms from a source population may be considered as sequences of discrete symbols generated by a ‘source’ distribution, where the capacity to sequence long stretches of genomes is congruent with the use of large block sizes in the design of communication channels. Rather than arising *temporally as an ordered sequence of* symbols in a communication channel, genetic sequences are non-temporal linear outputs of a sequencing scheme. This perspective ultimately enables the utilization of important information-theoretic asymptotic properties in the analysis of population genetic data.

Specifically, we introduce the concept of *typical genotypes* for a population, analogous to the core notion of typical sequences in information theory. These are genotypes one typically expects to encounter in a given population and are likely to represent the population very well. We analyze these typical genotypes from various perspectives. We show that it is possible that a genotype is typical to two different populations at once and give an algorithm that can quickly decide whether mutual typicality occurs, given standard models for two populations.

Crucially, we identify conditions in which it is *likely* that mutual typicality occurs asymptotically, that is, for genotypes consisting of a very high number of variants. What we observe, however, is that in this case, only a very small portion of typical genotypes for the latter population is typical for the first. This immediately suggests a classification scheme based on typical sets. We introduce two of such typical-set based classifiers and show that their error rates decay exponentially fast, as one would expect from a good classifier. Moreover, we show that such classifiers generally perform well even in the presence of sampling noise arising from small training sets.

From a mathematical point of view, a recurring difficulty is the non-stationarity of the source distribution, or in other words, that the markers vary in their frequency across loci. This prevents us from directly utilizing some of the standard results in information theory that apply to stationary sources, and required us to find more refined mathematical arguments instead.

## 1.1 Typical sequences and the *asymptotic equipartition property*

Information Theory (historically, Communication Theory) is at core concerned with the transmission of messages through a noisy channel as efficiently and reliably as possible. This primarily involves two themes, data *compression* (aka, *source coding*) and error correction (aka, *channel coding*). The former theme is mainly concerned with the attainable limits to data compression, while the latter involves the limits of information transfer rate for a particular source distribution and channel noise level. Both themes rely intrinsically on the notion of ‘typical sequences’.

A key insight of Shannon, the *asymptotic equipartition property* (AEP) forms the basis of many of the proofs in information theory. The property can be roughly paraphrased as “Almost everything is almost equally probable”, and is essentially based on the law of large numbers with respect to long sequences from a random source. Stated as a limit, for any sequence of i.i.d. random variables *X*_*i*_ distributed according to *X* we have,

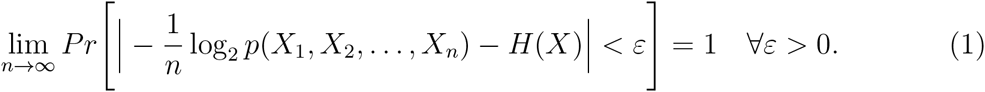

This property is expressed in terms of the information-theoretic notion of *empirical entropy*. This denotes the negative normalized log probability of a sequence *x*, an entity better suited for analysis than *p*(*x*). This property leads naturally to the idea of typical sequences, which has its origins in Shannon’s original ground-breaking 1948 paper [Shannon, 1948]. This notion forms the heart of the central insights of Shannon with respect to the possibility of reliable signal communication, and features in the actual theorems and their formal proofs. The definition of a typical set 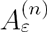 with respect a distribution source *X*, its entropy *H*(*X*), a (small) *ε* > 0 and a (large) *n*, entails the set of all sequences of length n that may be generated by *X* such that,

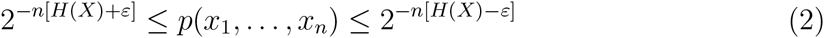

where *p*(*x*_1_, *x*_2_,…, *x*_*n*_) denotes the probability of any particular sequence from *X.*

If the source is binary and stationary it is intuitive to spot sequences that are possibly typical. For instance, say we have a binary independent and identically distributed (i.i.d) source with a probability for “1” of 0.1, then the sequence 0000100010000000000000100000 0011 seems very possibly typical (as it has roughly 10% 1s), while the sequence 0110100110 1100101111101001001011 is most probably not. Note that typical sequences are not the most probable ones; evidently, the most probable for this source is 0000000000000000000000000000000.

The interesting and useful properties of typical sets are a result of the AEP, and are thus *asymptotic* in nature: they obtain for large enough *n*, given any small arbitrary ’threshold’ *ε*. Formally, for any *ε* > 0 arbitrarily small, *n* can be chosen sufficiently large such that:

a. the probability of a sequence from *X* being drawn from 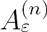 is greater than 1 – *ε*, and
b. 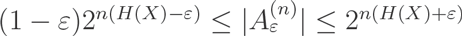

Thus at high dimensionality (*n* ≫ 1), the typical set has probability nearly 1, the number of elements in the typical set is nearly 2^*nH*(*X*)^, and consequently all elements of the typical set are nearly equiprobable with a probability tending to *2*^*−nH*(*X*)^ ([Cover and Thomas, 2006] Theorem 3.1.2).

The set of all sequences of length *n* is then commonly divided into two sets, the *typical set*, where the *sample entropy* or the *empirical entropy*, denoting the negative normalized log probability of a sequence, is in close proximity (*ε*) to the true entropy of the source per Eq. (2), and the non-typical set, which contains the other sequences (Fig. 1). We shall focus our attention on the typical set and any property that is true in high probability for typical sequences will determine the behaviour of almost any long sequence sampled from the distribution.

**Fig. 1:**
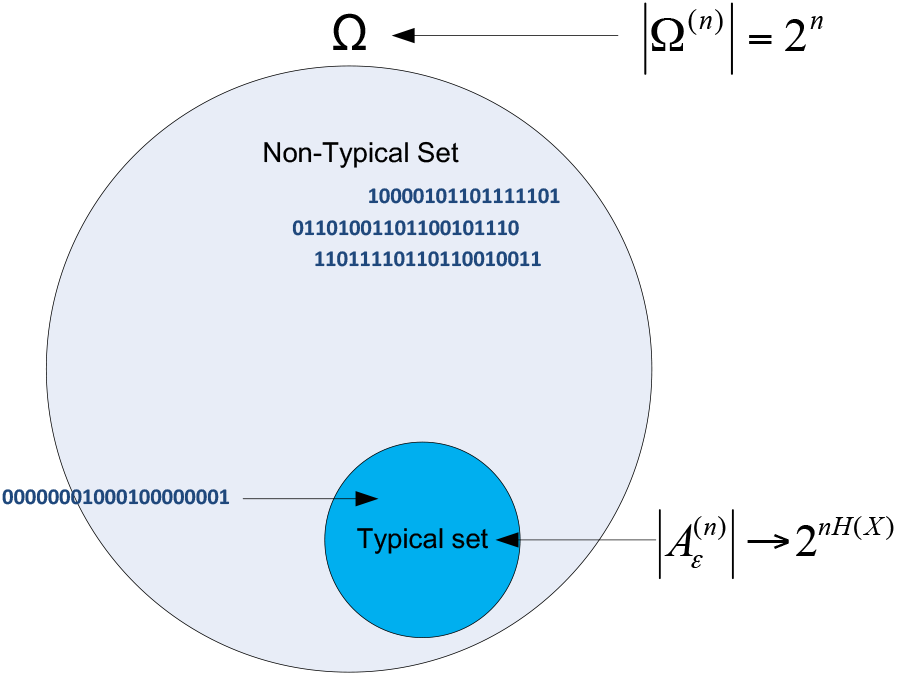
The universe of all possible sequences with respect to a source distribution in a high dimensional space can be divided into two exclusive subsets, typical and non-typical. Here, we illustrate one typical sequence and a few very non-typical sequences corresponding to an i.i.d. source with probability of 0.1 for “1” for some small epsilon and high *n*.

## 1.2 The Population Model

We consider for simplicity two *haploid* populations *P* and *Q* that are in linkage equilibrium (LE) across loci, and where genotypes constitute in a sequence of *Single Nucleotide Polymorphisms* (SNPs). A SNP is the most common type of genetic variant – a single base pair mutation at a specific locus usually consisting of two alleles (the rare/minor allele frequency is >1%). Each SNP *X*_*i*_ is coded 0 or 1 arbitrarily, and SNPs from population *P* have frequencies (probability that *X*_*i*_ = 1) *p*_*i*_ while those from population *Q* have frequencies *q*_*i*_. Closely following practical settings, we assume some arbitrary small cut-off frequency for SNP frequencies, such that frequencies in any population cannot be arbitrarily close to fixation, 0 < *δ* < *p*_*i*_,*q*_*i*_ < 1 – *δ*. Each genotype population sample is essentially a long sequence of biallelic SNPs, e.g., GCGCCGGGCGCCGGCGCGGGGG, which is then binary coded according to the convention above, e.g., 0101100010110010100000. The probability of such a genotype *x* = (*x*_1_, *x*_2_,…,*x*_*n*_) from *P* is then *p*(*x*) = (1 – *p*_1_)*p*_2_(1 – *p*_3_)*p*_4_*p*_5_…*p*_*n*_. We first assume the SNP frequencies are fully known (as if an infinite population sample is used in the learning stage), and later on relax this assumption in the section on small-sample related noise. Finally, for analyzing properties in expectation and deriving asymptotic statements we assume *p*_*i*_ and *q*_*i*_ are sampled i.i.d. from frequency distributions. For making explicit calculations and numerical simulations we employ a parameterized Beta distribution for SNP frequencies, such that *p*_*i*_ ~ *B*(*α*_*P*_, *β*_*P*_),*q*_*i*_ ~ *B*(*α*_*q*_, *β*_*q*_), as is standard in population genetic analysis ([Rannala and Mountain, 1997]). The use of a common Beta model for allele frequencies was adopted for both its mathematical simplicity and goodness of fit to empirical distributions from natural populations, and is by no means a prerequisite for arriving at our main results. Finally, to simulate our results, we sample SNP frequencies from these distributions and then sample long genotypes from the multivariate Bernoulli distribution for populations *P* and *Q* that are parameterized by *p*_*i*_ and *q*_*i*_, *i* : 1…*n*, respectively.

## 1.3 Properties of sequences of genetic variants

Population SNP data have several interesting ‘set-typicality’ properties that may render them amenable to information theoretic analysis:

a. SNPs typically are bi-valued, simplifying modeling SNPs as sequences of binary symbols from a communication source.
b. The standard assumption of *linkage equilibrium* within local populations translates to a statistical independence of *X*_*i*_, which in turn enables the applicability of the AEP (for a non-stationary source with independent symbols).
c. SNPs have typically differing frequencies across loci (i.e., analogous to a ‘nonstationary’ source), resulting in statistical terms in deviations from i.i.d. samples; this property makes an information theoretic analysis of SNP genotypes more challenging, being highly dependent on the existence of advanced forms of the AEP.
d. The recent availability of very large number of SNPs from high-throughput sequencing of genomes enables the consideration of very long sequences (size *n*), or ‘block sizes’ in information theoretic terms, with asymptotic qualities.
e. SNP frequencies are commonly above some arbitrary *cut-off frequency*, so that the *variance of* log_2_(*p*_*i*_) is bounded, a requirement for a nonstationary form of the AEP to hold (as we shall see).
f. SNPs typically have low minor allele frequencies (MAF) in natural populations (Fig. 2A). If we consider long sequences of SNPs as our genotypes, then the set of typical sequences from a population will be small (of asymptotic size 2^*nH*(*X*)^) relative to the ‘universe’ set (of size 2^*n*^) of all possible genotypes. This property enables treating such typical sequences as effective proxies for their source population.
g. Different populations often have different SNP-based genetic diversities (see the wide variation in heterozygosities across human populations in Fig. 2C), and SNP frequencies are often highly correlated between close populations (Fig. 2B). These properties have particular interpretations when populations are seen as communication ‘sources’.

**Fig. 2:**
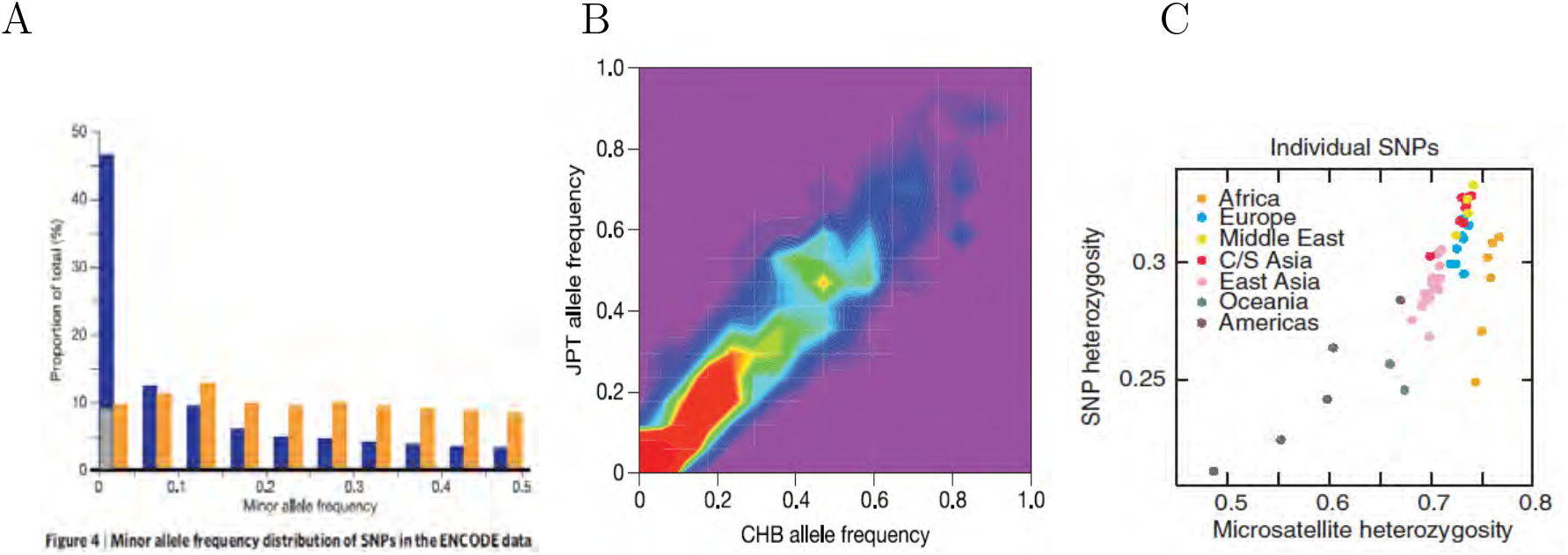
Human populations typically exhibit predominately low SNP frequencies (and thus commonly modeled by a Beta distribution highly skewed to the left), which are correlated between close populations (due to a split from common ancestry), and of differing average frequencies across worldwide populations. A: SNPs from the HapMap ENCODE regions according to minor allele frequency (in blue) [Borrowed with permission from *Nature 2005; 437(7063): 1299–1320*, Fig. 4]. | B: SNP frequencies from the HapMap ENCODE project between (the relatively close) JPT and CHB populations are highly correlated between the two populations at each locus [Borrowed with permission from *Nature 2005; 437(7063): 1299–1320*, Fig. 6]. | C: Differing SNP heterozygosity across worldwide populations with most diversity occurring in Africa and least in the Americas and Oceania. [Borrowed with permission from *Nature Genetics 38, 1251 – 1260 (2006)*, Fig. 3].

## 1.4 AEPs for genotypes from multiple populations

To formulate AEP statements for genotypes comprised of long stretches of population variants, we first define two central concepts: population entropy rate and cross entropy rate. The entropy of a population with respect to a set of loci has been previously invoked in formulating measures of population diversity or differentiation with respect to a single locus ([Lewontin, 1995]). Since SNPs typically have differing frequencies across loci, translating in information theoretic parlance to ‘non-stationarity’ of the source, one cannot simply employ entropy *H* as a variation measure of a population. Instead, we need to define a population *entropy rate* across loci. Thus, with respect to a set of SNP frequencies in population *P*,

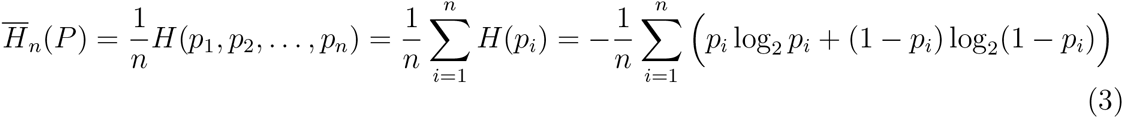

with the second equality due to independence across loci (absence of LD).^1^ We may now extend this concept by incorporating a second population that serves as the source, while the log-probabilities remain with respect to the first. In information theoretic terms, the cross entropy *H*(*p*, *q*) measures the average number of bits required to compress symbols from a source distribution *P* if the coder is optimized for distribution *Q*, different than the true underlying distribution. For univariate variables, the cross entropy can be expressed in terms of the Kullback Leibler divergence (also known as *relative entropy*),

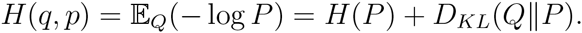

where we use lower-case in *H*(*p,q*) to distinguish this notion from the *joint entropy*, commonly denoted *H*(*P,Q*). The *population cross entropy* rate is then simply an average over *n* loci,

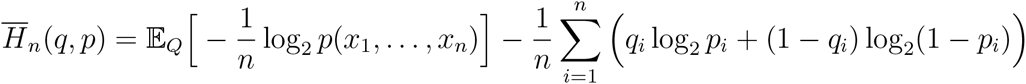

and similarly for 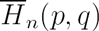.

Formally, if genotypes originate from distribution *P*, then by the non-stationary version of the AEP (see Appendix B.4.1 part 1) their log-probability with respect to *P* converges to the entropy rate of *P*,

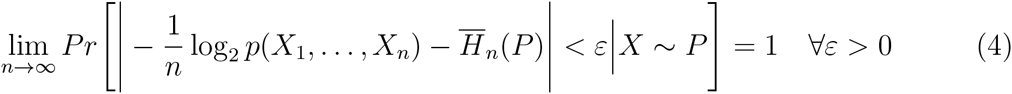

whereas if genotypes originate from distribution *Q*, then their log-probability with respect to *P* converges to the cross entropy rate of *Q* with respect to *P*, essentially a ‘cross entropy AEP’ for non-stationary sources (see Appendix B.4.1 part 2),

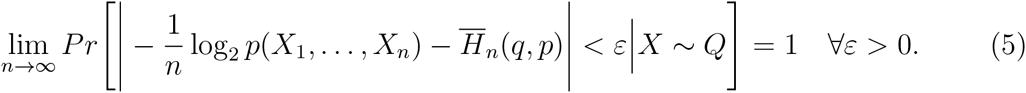

## 1.5 Typical genotypes

This consideration of the ‘set-typicality’ properties along with AEPs for our genotypes suggests that a notion of *typical-genotypes* may be fruitful for characterizing population samples. We therefore extend the standard definition of a typical set to support a non-stationary source, which better captures our population model. The set of typical genotypes of length *n* with respect to the *population entropy rate* of *P* and some small arbitrary *ε*, comprises of all genotypes whose frequency is within the bounds,

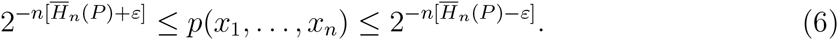

For notational simplicity, we will denote by *q*(*x*_1_, *x*_2_,…,*x*_*n*_) the corresponding probability of a genotype from population *Q*. Since the definition of a typical set pertains for any *n* and *ε*, our justification in invoking this concept in this context does not have to rely on asymptotic properties only, but holds naturally by virtue of commonly large *n* for SNPs.

## 1.6 Quantitative AEPs

It is beneficial to additionally formulate quantitative, non-stationary versions of the AEP theorems. Given that a genotype of length *n* is sampled from population *P*, the probability that it is not typical is bounded by

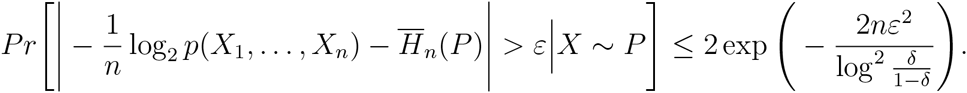

This estimate is proved in Appendix C.1. In the same way, the probability that the log probability under *P* deviates more than *ε* from the cross entropy rate, is estimated in the following quantitative version of a ‘cross entropy AEP’ for non-stationary sources,

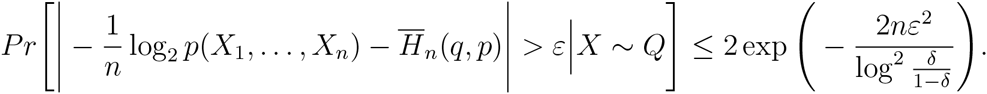

The corresponding non-quantitative versions of the AEPs in Eq. (4) and (5) are obtained by letting *n* approach infinity.

Since the above inequalities hold for every *n* and *ε >* 0, we can for instance choose,

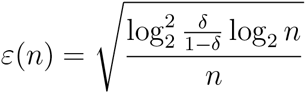

to conclude that,

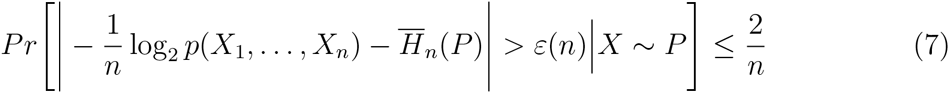

and similarly,

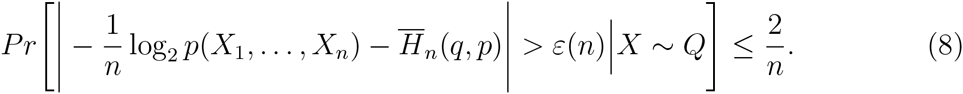

This shows that the deviation from the entropy rate practically scales as 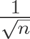, which is what one would expect also from a central limit theorem. A more careful analysis in Appendix C.1 also shows that the scale log^2^ 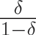 may actually be replaced by the sum

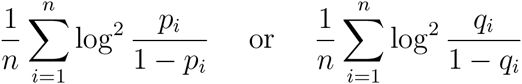

which for large *n* will be close to their expectation value and therefore are usually smaller for larger entropy rates. This may explain why the spread away from the entropy rate seems smaller for higher entropy rates. Fig. 3 depicts numerical simulations of the convergence rate of the AEPs under typical population scenarios.

**Fig. 3:**
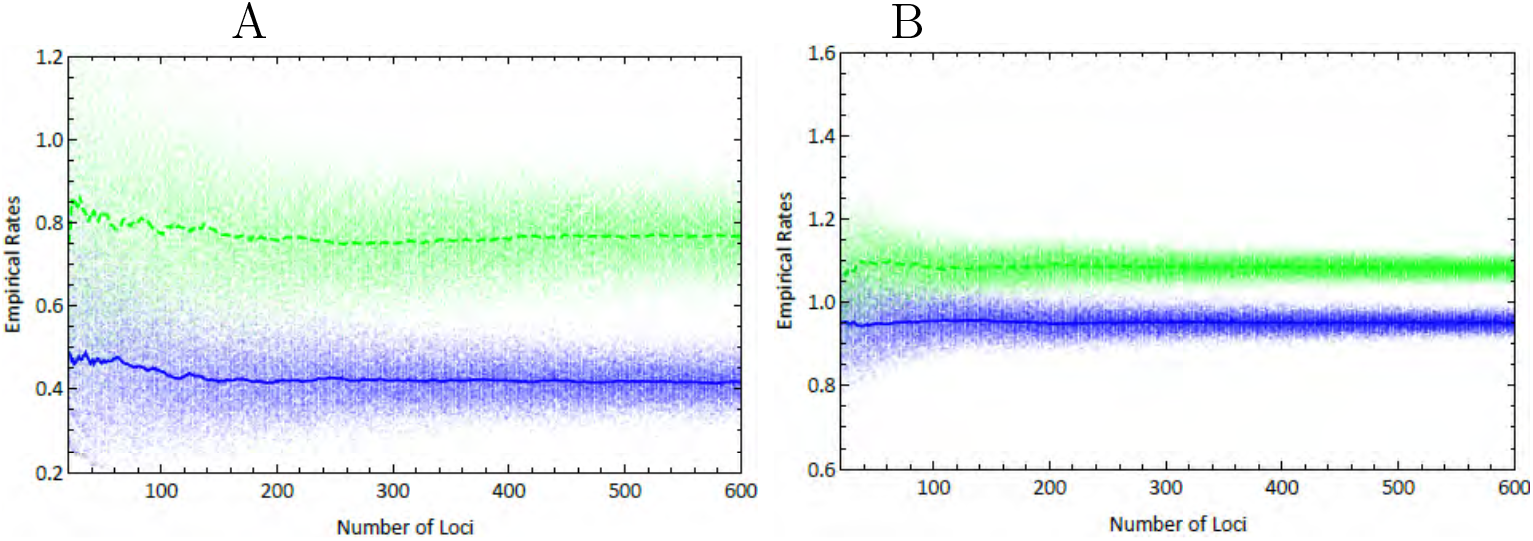
Numerical simulation of the convergence rate of the AEPs under two scenarios of population parameters, around the entropy rate 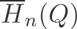 (blue) and the cross entropy rate 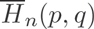 (green, dashed). A: Low entropy populations (Beta model w/*α*_*P*_ = 4/*β*_*P*_ = 20, *α*_*Q*_ = 2/*β*_*Q*_ = 20; *F*_*ST*_ = 0.032). | B: high entropy populations (Beta model w/ *α*_*P*_ = 24/*β*_*P*_ = 20, *α*_*Q*_ = 14/*β*_*Q*_ = 20; *F*_*ST*_ = 0.032).

## 1.7 The log-probability space

The AEP theorems of Eqs. (4-8) manifest as increasingly dense clusters of population samples on a log-probability space, centered on entropy and cross entropy rates, depending on their population of origin. To fully capture the interplay of genotype samples from the two source populations, and the information theoretic quantities of entropy and cross entropy rates, we take a two-dimensional perspective of the log-probability space. We should expect samples from the two populations to cluster around the *intersection* of the entropy and cross entropy rates of their respective populations, with a concentration that increases with the number of loci included in analysis. Crucially, typical genotypes should cluster tighter than general samples around the entropy and cross entropy rates intersection, since typical sequences are by definition constrained by some *ε* > 0. These results are illustrated in Fig. 4.

**Fig. 4:**
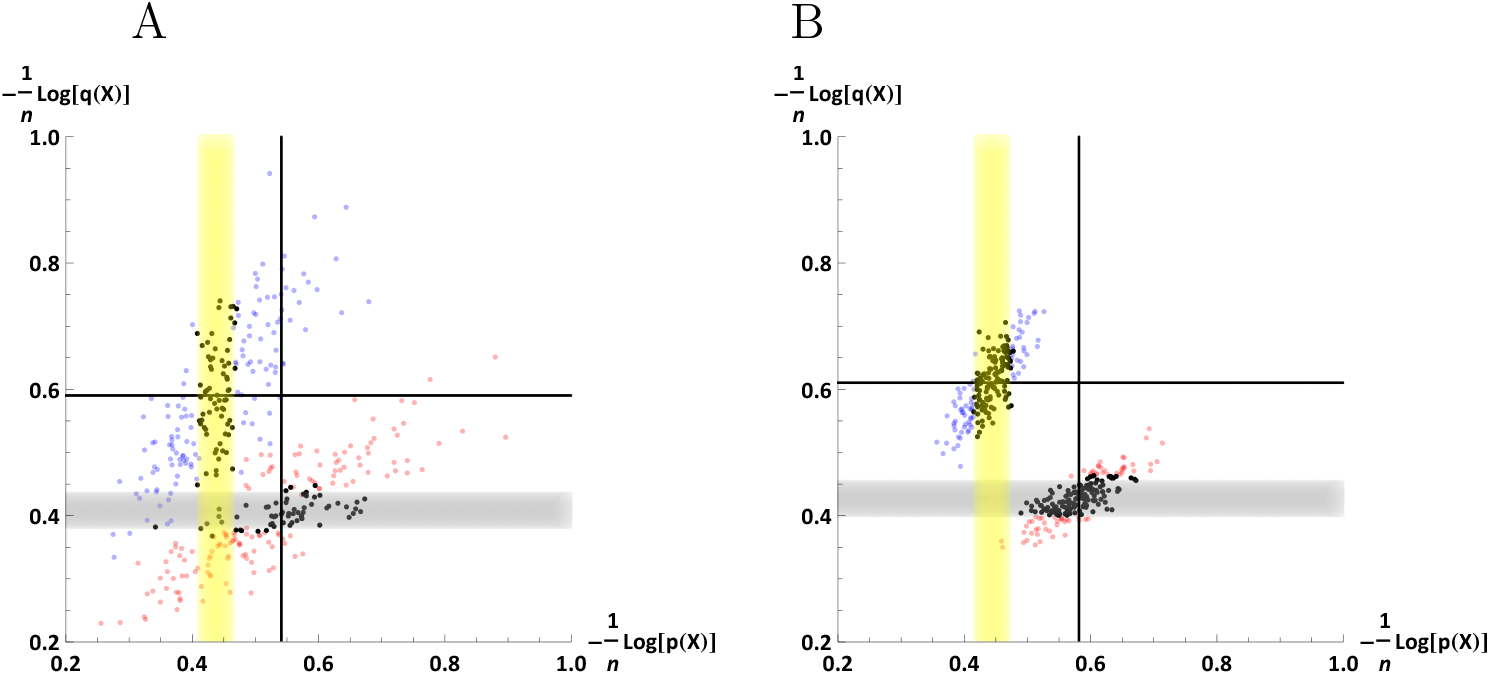
Samples from two different populations become clearly distinguished on a 2D log-probability plot when high number of loci are included in analysis, clustering around the intersection of the entropy (wide lines) and cross entropy (thin lines) rates of their respective populations. The width of the entropy stripes is twice *ε* to reflect the typicality criteria of Eq. (6), where here *ε* = 0.03. In this simulation, 200 genotype samples of 100 SNP loci (panel A) and 600 SNP loci (panel B) were drawn from each of the two populations of similar entropy rates and *F*_*ST*_ = 0.05, where allele frequencies were modeled on Beta distributions (*α* = 1, *β* = 8 for both populations).

The divergent modes of concentration on the log-probability plot of samples from the two populations suggest that the *proximity* of the entropy and cross entropy rates is an important metric in the context of population assignment for genotypic samples, as we shall see in what follows.

## 2 Set perspective on typical genotypes

Before we approach the task of constructing classifiers for population genetic samples based on the notion of typicality, we present two perspectives on the interplay of typical sets: from their set-overlap and exclusivity, and from their geometric dispersion. In particular, we will be interested in the asymptotic properties due to the high dimensional nature of genotypes (with the inclusion of large number of SNPs). Our hope would be that under expected population model of real population SNP data, sets of typical genotypes from diverse populations *asymptotically* become non-overlapping and good proxies for their respective sources.

### 2.1 Mutual and exclusive typicality

We first define the concept of mutual typicality. Formally, given *P, Q* and small *ε*_*p*_ > 0 and *ε*_*q*_ > 0, we would like to know whether the two typical sets partially overlap, i.e., is there at least one *x* = (*x*_1_,…,*x*_*n*_) such that *x* is *mutually typical* to both *P* and *Q?* Any such sequence *x* would need to satisfy the two inequalities,

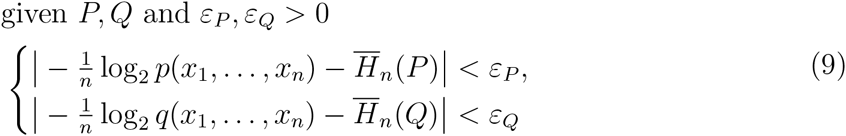

or equivalently as a set of four *linear programming* inequalities of degree *n*,

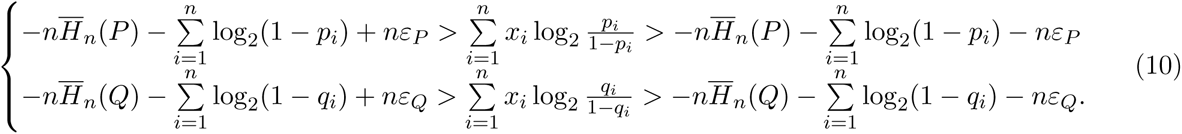

Notice that our notion of mutual typicality is conceptually different to the standard information theoretic concept of ‘joint typicality’, which concerns whether two different sequences are each marginally typical and at the same time typical with respect to the joint distribution (a central concept in Shannon’s channel coding theorem).

The above formulation (for a finite *n*) is essentially a 0 – 1 *integer programming with no optimization* problem: given *n* Boolean variables and *m* (= 4 in this case) linear constraints, the problem is to find an assignment of either 0 or 1 to the variables such that all constraints are satisfied ([Impagliazzo et al., 2014]). The ‘no optimization’ qualification reflects the omission of an objective function to be optimized that is usually an integral part of a linear programming framework, while only considering the problem of deciding if a set of constraints is feasible. This special case of an integer programming is a decision rather than optimization problem, and as such is *NP–complete* rather than *NP–hard*. In fact, 0–1 *integer programming with no optimization* is one of *Karp’s 21 NP–complete problems* ([Zuckerman, 1996]). Crucially for our purposes, the NP completeness means that it is not readily amenable to resolution for a large *n*, as our genotypic framework typically demands. Nevertheless, for small values of *n* one may solve the integer programming problem and infer the existence of mutual or exclusive typicality.

As with other NP-complete problems, high-dimensional instances are intractable and so heuristic methods must be used instead. We shall see that for large *n*, an approximate solution to the problem of mutual typicality can be found very efficiently, since the integer programming problem is well approximated by a *linear* programming problem. We slightly simplify the problem, making it effectively independent of the choice of *ε*_*P*_ and *ε*_*Q*_. Thus, we ask whether given *any* small *ε*_*P*_ and *ε*_*Q*_ there exists an overlap of the two typical sets for high values of *n*. Next, we simulate the log-probability space with samples drawn from a *uniform* (i.e., max entropy) distribution, so that a maximal set of different genotypes from the total *2*^*n*^ universe is captured. The cross entropy AEP of Eq. (5) directly implies that asymptotically the density of this domain is concentrated at the intersection of two cross entropy rates, 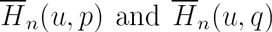, where *U* is the uniform distribution. This coordinate may be expressed as a function of the SNP frequencies of *P* and *Q*,

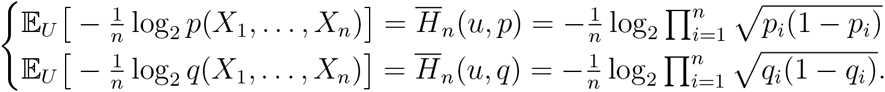

The contour of this domain is prescribed within boundaries which are the maximal and minimal empirical entropy values with respect to *P* and *Q* for any of the possible *2*^*n*^ genotypes,

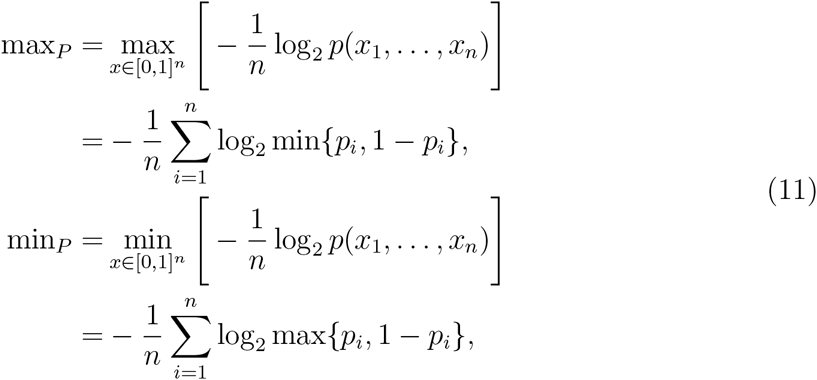

and similarly for population *Q*.

From Eq. (11) it is evident that these boundaries are an *average* across loci and therefore will depend on the parameters of the population model, rather than on the dimensionality *n*. However, since the domain inscribed by all possible samples on the log-probability space does not include the whole rectangular area prescribed by the boundaries, knowledge of these boundaries is insufficient for determining whether the intersection of the two entropy rates (i.e., the location where samples are asymptotically mutually typical) lies within the domain or is external to it.

In Theorem C.3.1 in the appendix we actually show that the domain converges (in the so-called Hausdorff distance) to a fixed, convex set, and provide an expression for the *contour* of this domain. The converge rate is approximately 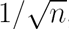, and therefore even for relatively small values of n the convex set is already a good approximation for the domain. This formulation, in conjunction with the entropy rates of *P* and *Q*, will then allow to immediately determine whether asymptotically there are mutually-typical genotypes (a solution to Eq. (10) for high *n*): if the intersection of the two entropy rates lies within the genotype domain then for any *ε*_*P*_ and *ε*_*Q*_ chosen as small as we wish, there will be mutual typicality for some non-empty subset of genotypes; else, there will only be exclusive typicality (a consequence of the convergence in the Hausdorff distance at the given rate is that the domain is sufficiently non-porous, with porousness bounded by 1/*n*). Fig. 5 depicts numerical simulations of this domain along with its computed contour at the asymptotic limit, for two representative scenarios of mutual and exclusive typicality.

**Fig. 5:**
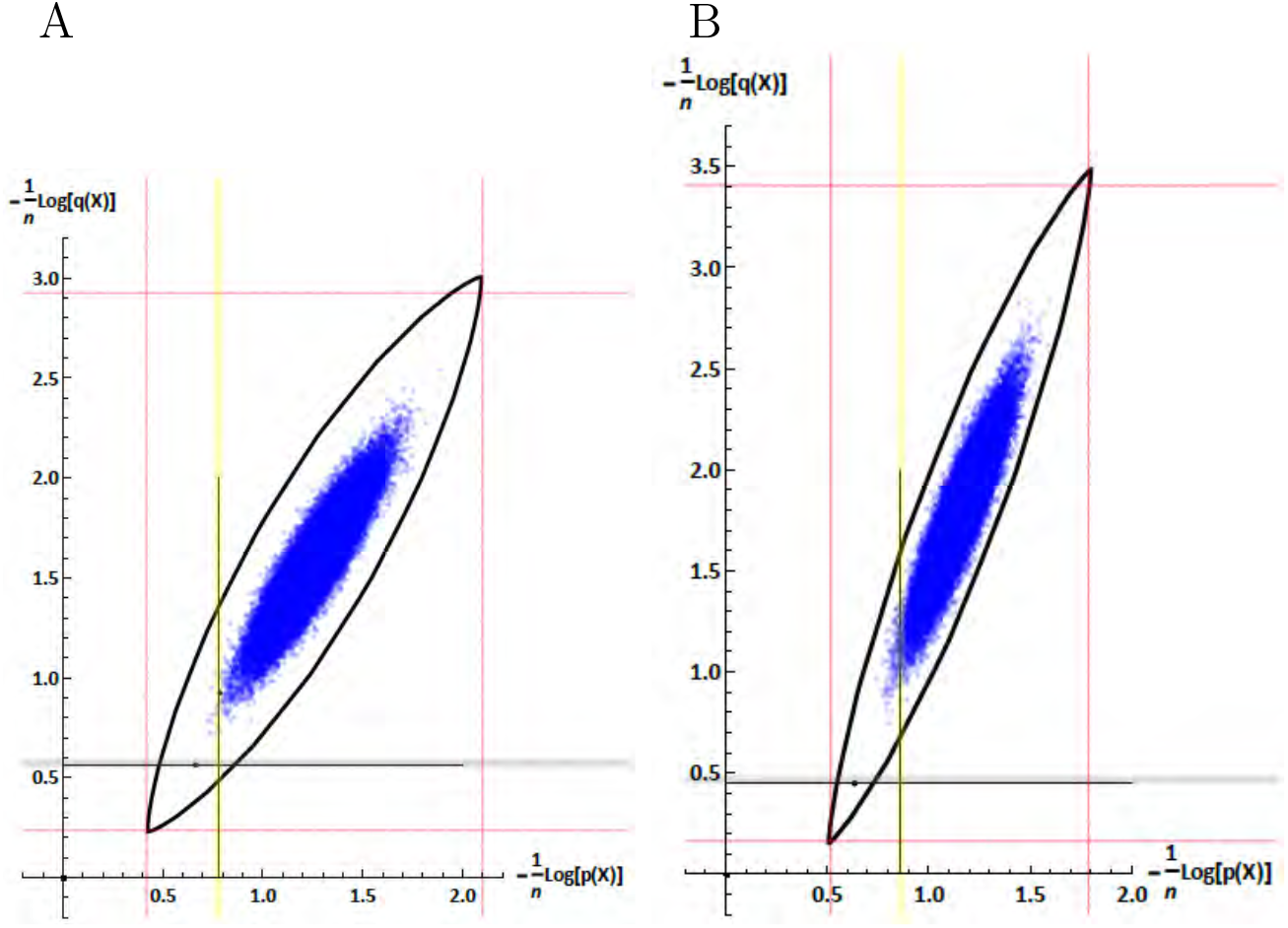
Instances of ‘source-less’ mutual and exclusive typicality scenarios for populations *P* and *Q* at the asymptotic limit for n. A simulation of the analytic formulation of a contour of the domain inscribed by all samples drawn from the uniform distribution over the space, was overlaid on top of a simulation of a plot of samples from this uniform distribution, with respect to their log-probability. The wide stripes represent the entropy rates of *P* (yellow) and *Q* (grey). The thin border lines represent the minimum and maximum attainable values for samples from the specific population distributions. A: the intersection of the two entropy rates lies within the domain, implying existence of mutual typicality (populations modeled on Beta distributions for SNP frequencies with *α*_*P*_ = 6/*β*_*P*_ = 18; *α*_*Q*_ = 3/*β*_*Q*_ = 18, and using *n* = 40 loci and 60K samples in the domain simulation). | B: the intersection lies outside the domain, implying merely exclusive typicality (populations modeled on Beta distributions for SNP frequencies with *α*_*P*_ = 15/*β*_*P*_ = 36; *α*_*Q*_ = 4/*β*_*Q*_ = 36, and using *n* = 40 loci and 60K samples in the domain simulation). The intersection of the cross entropy and entropy rates are marked as small dots on the entropy rate lines, merely to indicate where highest density would be if genotypes were sampled from *P* and *Q*, rather than from the maximum entropy distribution.

From a set perspective, this result translates into two scenarios for the interplay of typical sets at the asymptotic limit: [a] if the intersection of the entropy rates lies within the contour of the log-probability domain then the two typical sets will have some overlap, whereas [b] if the intersection lies outside the contour then the two typical sets will completely separate. Since we assume arbitrarily small *ε*_*P*_ and *ε*_*Q*_, the set overlap in case [a] only depends on the density of the domain at the intersection of the entropy rates, and is approximately given by 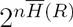, where *R* is the distribution given by frequencies *r*_*i*_ that yields the maximum entropy rate under the constraints that 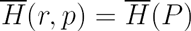 and 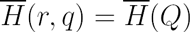.

To see that there could not be a third scenario in which one typical set is wholly contained in the other (except trivially for the hypothetical case where one distribution is uniform, i.e., 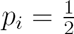), we show that the entropy rate cannot coincide with the minimal or maximal bounds of the domain on the log-probability space. From a geometric perspective on the log-probability space (see Fig. 5) this means that the two entropy rate lines are never tangential to the genotype domain. Formally, with respect to the minimum for population *P* from Eq. (11), the inequality,

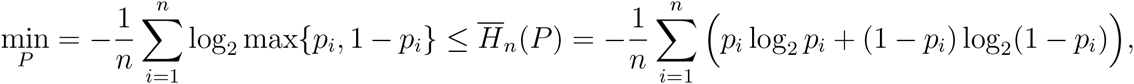

obtains equality only for *p*_*i*_ = 1/2 for all *i* : 1…*n*, an impossible population scenario (similarly for max_*P*_, min_*Q*_ and max_*Q*_). Fig. 6A depicts these possibilities in the form of Venn diagrams.

**Fig. 6:**
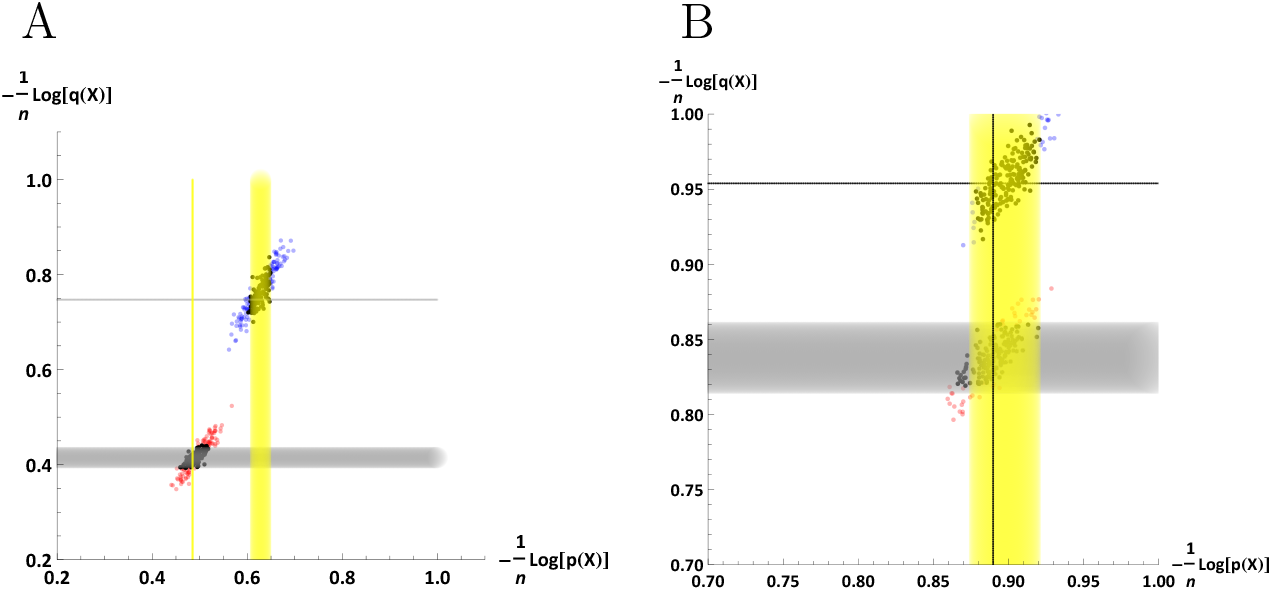
With samples originating from populations *P* and *Q*, there is *with probability* 1 either exclusivity of typicality (A) or complete one-sided mutual typicality (B). Entropy rates are marked as wide strips according to respective epsilons and cross entropy rates are the thin lines. A: a typical scenario in which there is exclusivity of typicality (*F*_*ST*_ = 0.02, *n* = 1000, *ε*_*P*_ = *ε*_*Q*_ = 0.02). | B: a highly uncommon scenario where one *cross entropy criterion* is close to zero although populations are distant (*F*_*ST*_ = 0.02, *n* = 1600, *ε*_*P*_ = *ε*_*Q*_ = 0.02), and therefore all samples from *Q* are mutually typical but none of *P* are as such (a zoomed view to capture the proximity of the entropy rate and cross entropy rate for *P*, the latter accentuated as black line).

### 2.2 Source-full mutual typicality

We would also like to analyze a modified definition of mutual typicality, which only considers probable genotypes, i.e., those likely to originate from their respective populations by a random sampling procedure. We also retain the original relevance of the choice of *ε*_*P*_ and *ε*_*Q*_, and again focus our inquiry at the asymptotic limit. This perspective on mutual typicality is explicitly pertinent for our subsequent inquiry into typicality-based classifiers. It is now necessary to introduce the concept of ‘cross entropy criterion’, which measures the proximity of the entropy and cross entropy rates. There are two such criteria for our two-population framework,

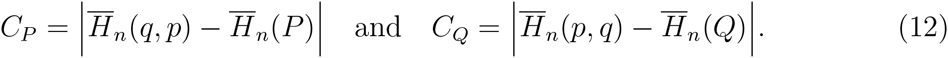

Clearly, if the two populations are effectively a single population (P=Q) then both cross entropy criteria will be zero, since from basic definitions,

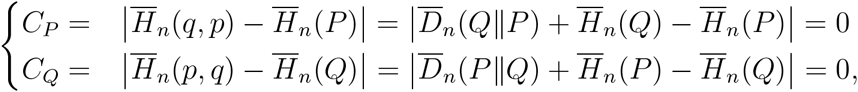

where the KL-Divergence rate from *P* to *Q* is naturally defined as,

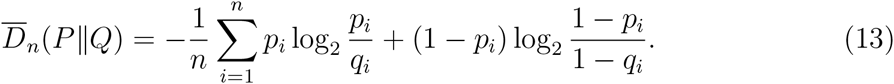

(and similarly from *Q* to *P*). However, one cross entropy criterion may be asymptotically zero under a standard model for allele frequencies, even given *differing* populations; population clusters are then inseparable on the corresponding log-probability plot along the corresponding axis (Appendix B.2). Crucially, both criteria cannot asymptotically be zero *at the same time* (Appendix B, Remark B.2.1),

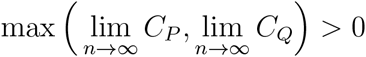

Now, from the AEP and the cross entropy AEP of Eqs. (4) and (5) it follows that the predominant asymptotic scenario is exclusive typicality *with probability* 1, given a choice of small typicality *ε’s* based on the *cross entropy criteria*, such that *ε*_*P*_ ≤ *C*_*P*_ and *ε*_*Q*_ ≤ *C*_*Q*_. Otherwise, in case *CP* < *ε*_*P*_ or *C*_*Q*_ < *ε*_*Q*_, then *asymptotically* one typical set will be *with probability* 1 fully contained in the other (i.e., all samples originating from one population are mutually typical and all samples originating from the other population are exclusively typical). These two cases are depicted in Fig. 6, under large *n* to simulate the asymptotic behavior.

Let 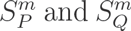 denote random samples of size *m* from population *P* and *Q* respectively. Define the sampled typical sets 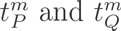 by,

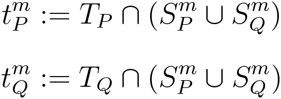

If the sample size *m* is not too large, the Venn diagram associated with these two sets is most likely equal to one of the two options depicted in Fig. 7B.

**Fig. 7:**
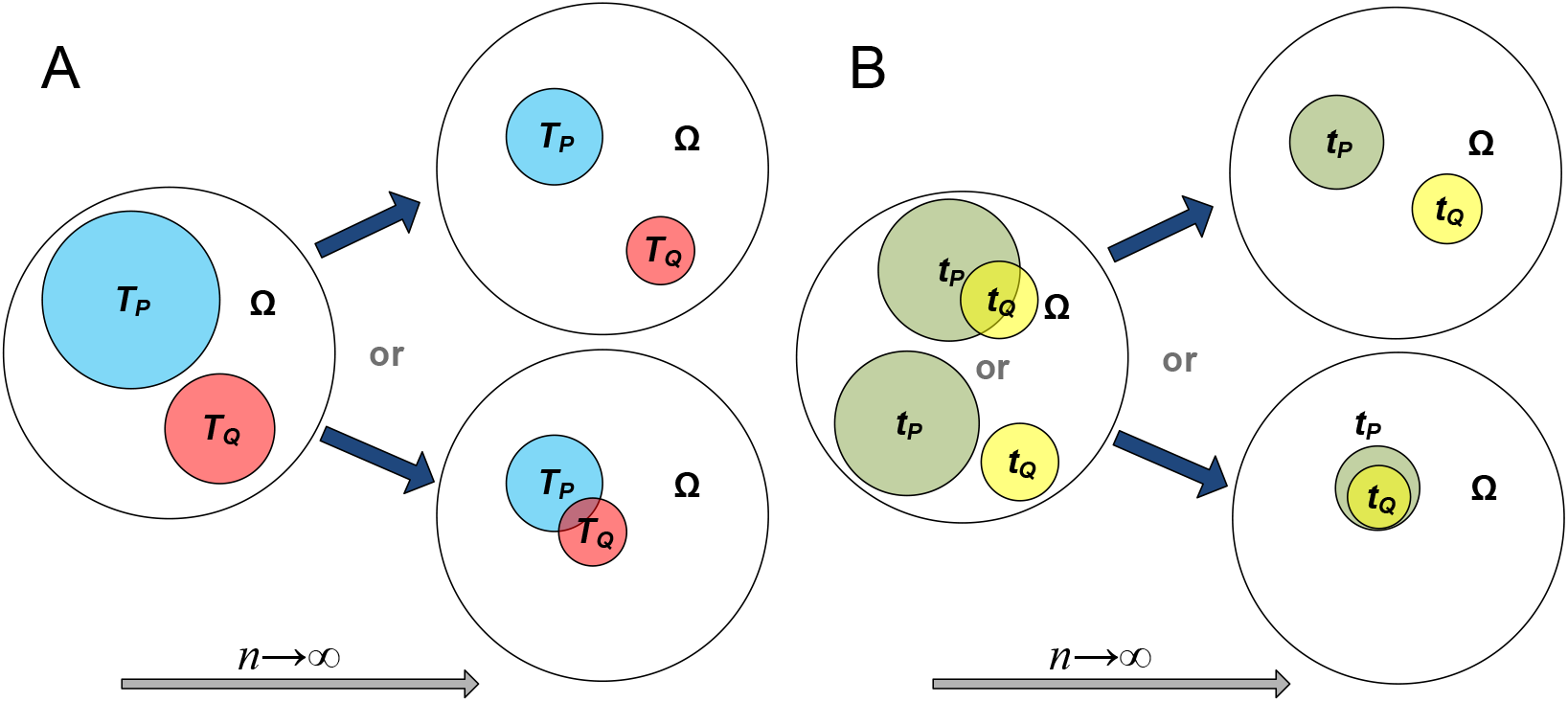
A Venn diagram of the interplay of two typical sets (denoted *T*_*P*_ and *T*_*Q*_) with respect to populations *P* and *Q*, from low n to an asymptotic limit. A: In the general case where we consider all possible genotypes from the universe, exclusive typicality at low dimensions transforms into either complete separation (bottom) or a very slight overlap (top), depending on the model parameters of the two populations. | B: In the case where genotypes are sampled from their source populations, a possible overlap in low dimensions transforms into either complete separation (top) or, rarely, a case where one typical set is wholly contained in the other (bottom). Note that the size of the typical sets relative to the universe is asymptotically zero, an aspect that that cannot be captured in this schematic.

## 3 A geometric perspective

We can gain more insight into the relation of typical genotypes to non-typical ones by taking a geometric perspective, where long genotype sequences are seen as vectors in *n*-dimensional genotype space [Huggins et al., 2007]. Essentially, the genotypes all lie on a subset of the vertices of a hypercube of dimension *n* (Fig. 8).

**Fig. 8:**
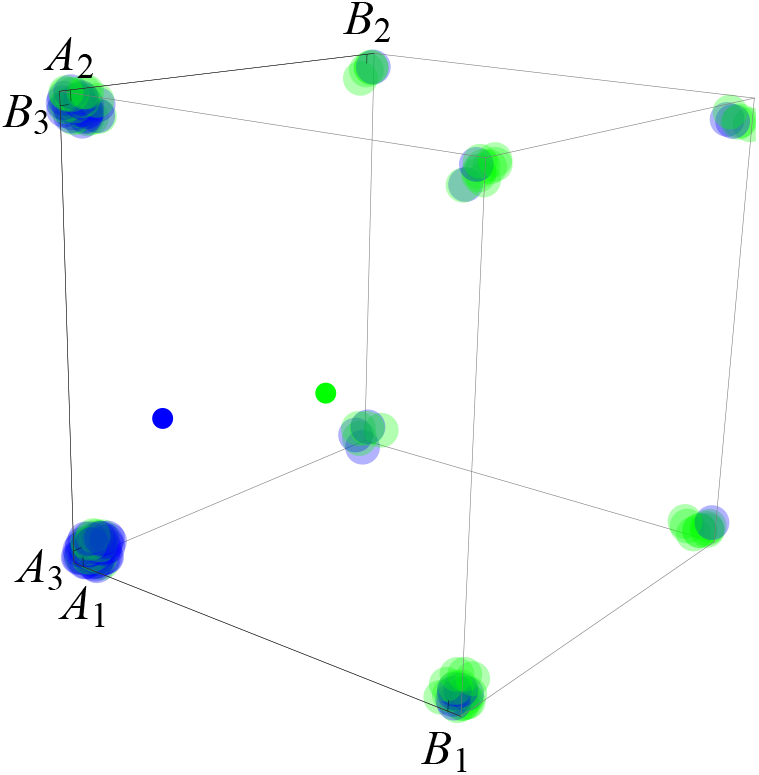
A geometric representation of the space of 3 SNP genotypes sampled from two populations. Genotype samples lie on the vertices of the (hyper)cube, where *A*_*i*_ is the “0” allele and *B*_*i*_ the “1” allele for locus *i, i* : 1…3 (e.g., genotype samples on the bottom left vertex *A*_1_*A*_2_*A*_3_ are 000 genotypes). Here 40 samples were drawn from one population (blue) and 40 samples from the other population (green), with respective population centroids represented by smaller dots within the cube.

How are the typical genotypes dispersed with respect to hypercube space? From the inequalities of Eqs. (10) it is evident that all typical genotypes are represented by those vertices that lie inside an (*n* – 1)-dimensional hyperplane of width 2*ε* intersecting the hypercube at some point, with an orientation and location fully determined by the parameters of the population distribution.

More importantly, at high dimensions the set of typical genotypes disperses evenly across the space occupied by population samples. The evidence for this comes from two types of numerical simulations. First, a PCA plots, which are known to essentially retain relative distances in the largest principal components, clearly indicate that typical genotypes behave as a random sample from the population, as depicted for two different populations in Fig. 9.

**Fig. 9:**
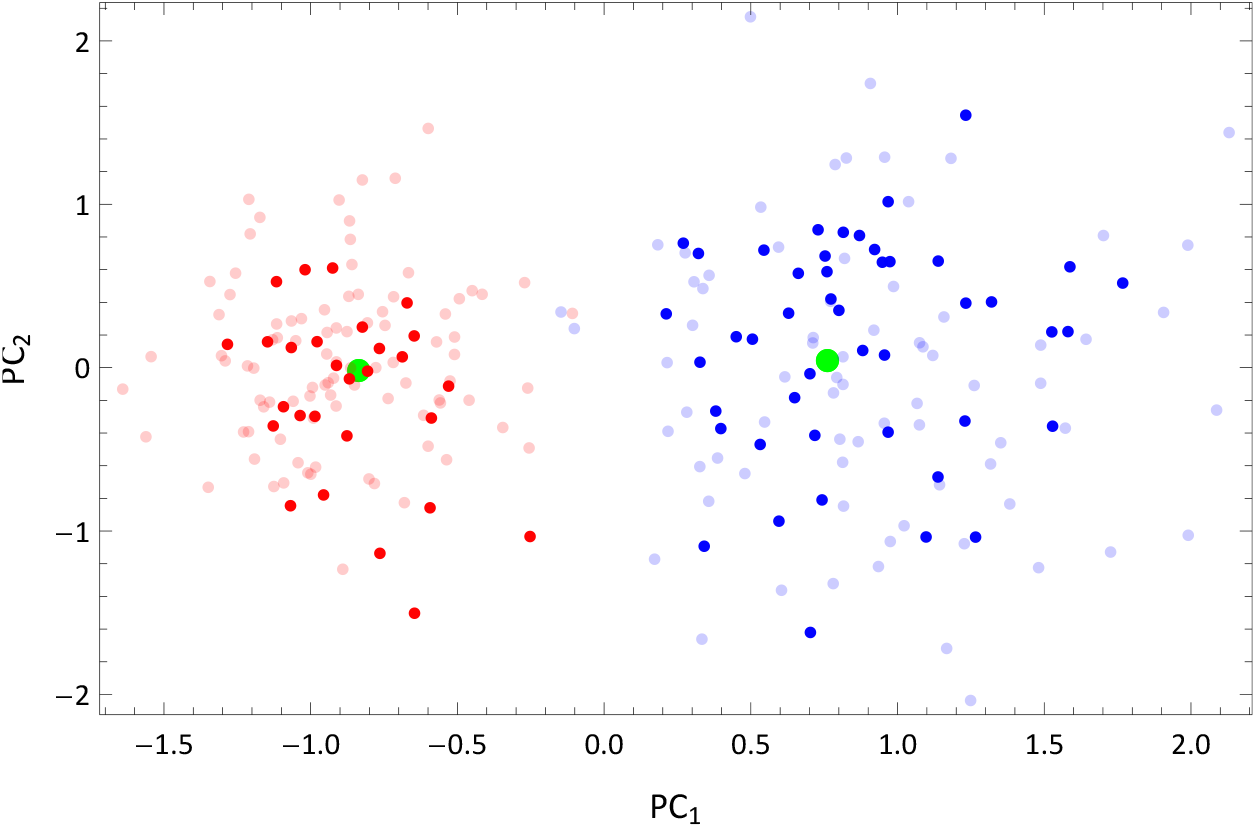
A PCA plot of two populations, blue and red, with typical genotypes of each in dark blue and dark red respectively (with centroids in green), demonstrating the even dispersion of typical samples in high dimensions. The simulation uses 120 samples of n=180 loci drawn from each population and SNP frequencies modeled on Beta distributions (*α*_*P*_ = 4, *β*_*P*_ = 20, *α*_*Q*_ = 2, *β*_*Q*_ = 20, *ɛ* = 0.01).

Second, an analysis of the average pairwise distance of typical genotype pairs compared to that of the whole distribution, reveals that the former converges to the latter *even when only a small portion of the pairs are typical* (see Appendix B.3 for the asymptotic equidistance property; see [Granot et al., 2016] for the effect of LD on equidistance). Note that trivially, if the whole sample becomes typical at some high dimension then the two averages will by definition converge to the same value. Moreover, simulations at low dimensions reveal that typical genotypes are slightly more densely clustered than samples from the whole population, since the convergence to the total average distance is always from below. These results are illustrated in Fig. 10.

**Fig. 10:**
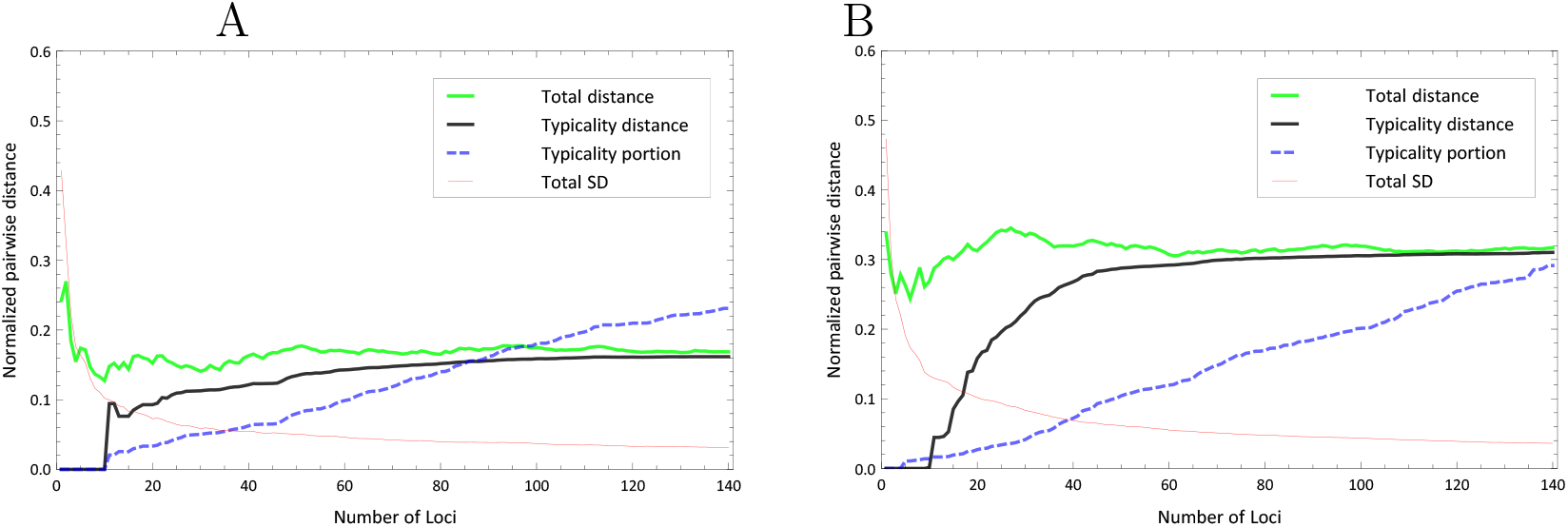
Two runs of a numerical simulation for average pairwise distance for samples drawn from a single population (in green), compared to a subset which comprises only of pairs of typical genotypes (in black), with *ε* = 0.01. The two curves always converge at high number of loci *n* even when only a small portion (in dashed blue) of the pairs are typical. We also convey the variance (thin red) of the pairwise total distance to highlight the asymptotic equidistance property. A: a scenario with population entropy rate = 0.41 (corresponding to very low MAFs)| B: entropy rate = 0.73 (corresponding to medium MAFs). Simulated using 120 samples drawn from a populations modeled on Beta distributions for SNP frequencies.

Not very surprisingly, the higher the population entropy rate the higher the average pairwise distance, since genotypes will tend to differ across more loci (see Appendix B.3). Finally, the lower the *ε* we choose to define our typical set the lower the rate of convergence: this suggests that genotypes which are essentially more ‘strongly typical’ (i.e., that correspond to a greater proximity to the entropy rate) are more tightly clustered.

## 4 Information-theoretic learning

The relation of information theory to statistical learning is currently a very active field of inquiry. The use of information theoretic learning criteria in advanced learning models such as neural networks and other adaptive systems have clearly demonstrated a number of advantages that arise due to the increased information content of these criteria relative to second-order statistics ([Erdogmus and Principe, 2006]). From a machine learning perspective, one of the early insights of information theory was to consider a classification problem as a noisy channel. Fano’s inequality ([Fano, 1961]), central to information theory, links the transmission error probability of a noisy communication channel to standard information theoretic quantities such as conditional entropy and mutual information.

We propose taking a further step in this direction, by implementing classifiers for genetic population data based on the principle and properties of typical sets, making use of our notions of population entropy rate, cross entropy rate, cross entropy criteria and typical genotypes. We derive our motivation by the preceding geometrical and mutual typicality analyses. The former perspective indicates that typical genotypes are asymptotically good representatives of their source populations, while the latter perspective indicates that samples from different populations are asymptotically *exclusively* typical. Crucially, we shall see that the performance of typicality-based classifiers is highly dependent on the value of the cross entropy criteria, specifically that,

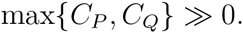

It is also instructive to compare the performance of such information-theoretic classifiers against a standard Bayes classifier (or *maximum-likelihood* classifier if no prior is available). This classifier is both conceptually simple in its definition, and optimal in its performance under known class-conditional densities. The expected error or misclassification rate of the Bayes classifier is called the *Bayes error* ([Hastie et al., 2009]). Our standard assumption of linkage equilibrium within populations (absence of within-class dependencies) motivates use of a *naïve Bayes* classifier, where class-conditional likelihoods are expressed as the product of allele frequencies across the independent loci.

### 4.1 Classifiers based on set-typicality

According to the AEP, if a *long* genotype is not typical for population *P*, then it is very unlikely that the genotype originated from population *P*. This suggests that a test of typicality could classify genotypes to the two different populations: naively, a genotype is classified to *P* if it is typical for *P*, and classified to *Q* if it is typical for *Q*. However, this naïve formulation of the classifier does not specify what should happen in case a genotype is typical to both *P* and *Q*, or if it is not typical to either population. Moreover, the definition of typicality is associated with a parameter *ε*. The choice of this parameter is closely related to these issues. Nonetheless, our previous analysis shows us how we may deal with these. Fig. 11 depicts a typical instance of the mapping of our population clusters on a 2D log-probability plot, in relation to the entropy and cross entropy rates, and some *ε* parameters.

**Fig. 11:**
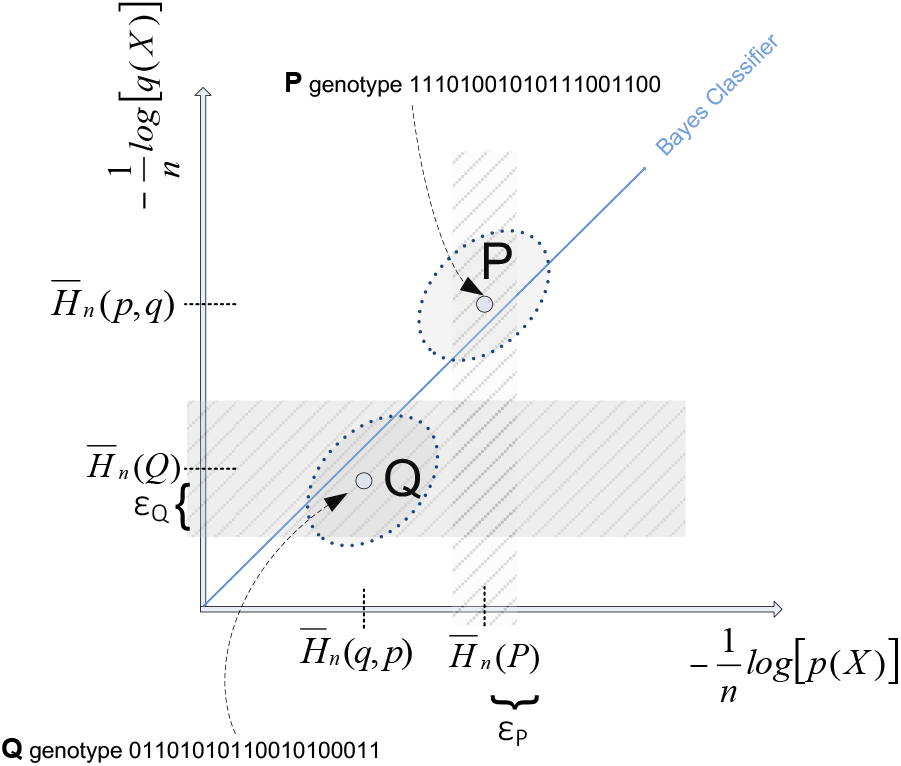
A typical instance of the location of the two population clusters on a 2D log-probability plot, in relation to the entropy and cross entropy rates, and a Bayes classifier (here 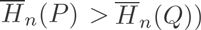. The centers of *P* and *Q* will *always* lie on opposite sides of the Bayes classifier diagonal since the KL-Divergence is always positive when populations differ (in terms of the coordinates of the two cluster centers, 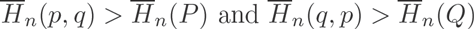.

We now introduce two typicality-based classifiers. To assess the performance of such a classifier, we estimate its error rates, which is the probability the classifier makes an error under the following process. With probability half, a genotype is sampled from population *P*, and with probability half, a genotype is sampled from population *Q*. Based on this genotype, the classifier guesses whether it originates from population *P* or from population *Q*. The error rate is the probability that the classifier guesses wrong. More precisely

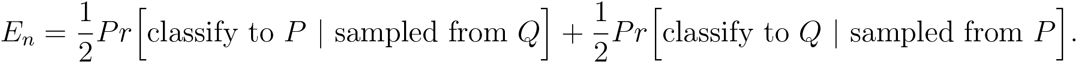

### 4.2 The naïve typicality classifier

The naïve typicality classifier is based on the idea of classification we have described before, that is classify to *P* (to *Q*) if the genotype is typical for population *P* (Q). As discussed before, we need to decide what the classifier should do when a genotype is typical for both populations. We prescribe that in this case of mutual typicality, the genotype will be classified to the population with the lower entropy rate, since the lower entropy rate population has higher asymptotic genotype probability, 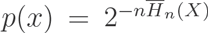 ([Cover and Thomas, 2006]). The classifier is then described by,

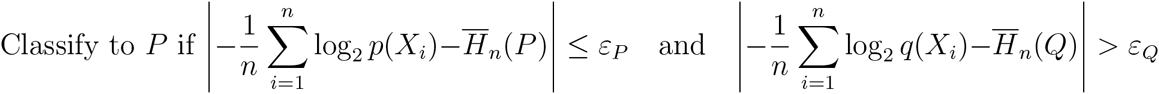

or else,

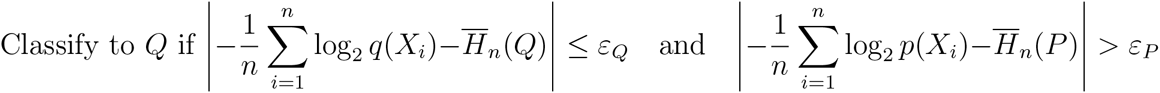

or else, if a genotype is not typical to any population, the classifier assigns by proximity, that is, it classifies to *P* if

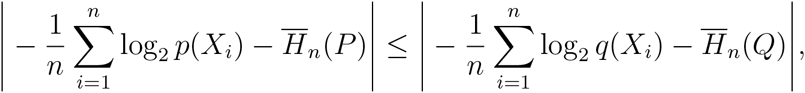

and otherwise to *Q*.

Or else, if mutually typical classify to *P* if, 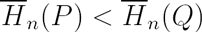, and otherwise to *Q.*

The choice of *ε* should not be arbitrary and also not necessarily equal between the two populations. If we choose *ε* too large we may never have exclusivity (as from some low dimension onwards all genotypes may be mutually typical), while if we choose *ε* too small we will not have typicality at lower dimensions (low SNP count). A reasonable choice is to base the two *ε*’s on the cross entropy criteria, which consequently have to be determined in the learning stage,

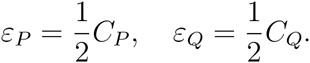

This represents a balance between avoiding mutual typicality (by setting *ε* not too high) while allowing for exclusive typicality (by setting *ε* not too low).

Based on the quantitative versions of the AEP and cross entropy AEP, we derive the following error bounds for the naïve typicality classifier (Appendix C.2),^2^

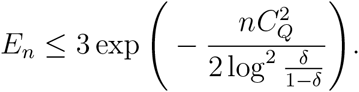

We note that a classifier which only classifies by proximity to the entropy rates amounts to the implicit assumption of equal entropy rates. This may lead to wrong classification of mutually typical samples, especially at lower dimensions; e.g., with differing entropy rates and with respect to the log-probability space, some samples from the cluster of *Q* may lie closer on the *x*-axis to 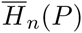 than on
the *y*-axis to 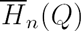, and thus be wrongly classified to *P*.

### 4.3 The cross entropy typicality classifier

In fact, our previous analysis of the cross entropy criteria shows that a simpler classifier, for which the selection of *ε* occurs implicitly and only one sample entropy is measured, would suffice. Without loss of generality, assume that *C*_*Q*_ > *C*_*P*_. Then classify to *Q* if the sample entropy with respect to *Q* of a genotype is closer to the entropy rate of *Q* than to the cross entropy rate of *P* given *Q*, i.e.,

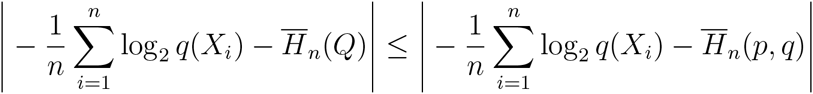

and classify to *P* otherwise.

Note that, without loss of generality, for any level of *C*_*Q*_, a higher convergence rate for our entropy and cross entropy AEPs implies that at any dimension *n*, samples from *Q* will tend to map tighter around 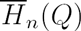, while samples from *P* will tend to map tighter around 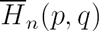 in the log-probability space. This immediately leads to stronger separation of the clusters along the *Q* axis, and therefore better classification prospects.

The error rate of this classifier can again be estimated from the quantitative AEPs, and is bounded by,^3^

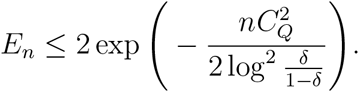

as shown in Appendix C.2.

The guiding principle behind this classifier is that the larger cross entropy criterion represents the empirical entropy dimension along which there is stronger separation between the clusters, a direct consequence of the AEP theorems of Eqs. (4) and (5). We note here that it is generally not possible for this classifier to avoid the computation of both *C*_*P*_ and *C*_*Q*_, inferring their relation by examining some simpler proxy.^4^. Indeed, the population entropy rates, which are generally more readily available, do not contain enough information since, for example,

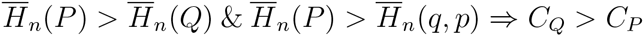

otherwise it is also possible that *C*_*Q*_ < *C*_*P*_ (Appendix B, Corollary B.2.2).

Specifically, if without loss of generality *C*_*Q*_ > *C*_*P*_ then the classifier considers the empirical entropy of samples from the two populations with respect to the *Q* distribution. For any given level of the cross entropy criterion (here *C*_*Q*_), a higher convergence rate roughly implies that at any dimension *n*, samples from *Q* will tend to map tighter around 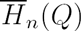, while samples from *P* will tend to map tighter around 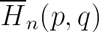. The two classifiers are presented schematically in Fig. 12.

**Fig. 12:**
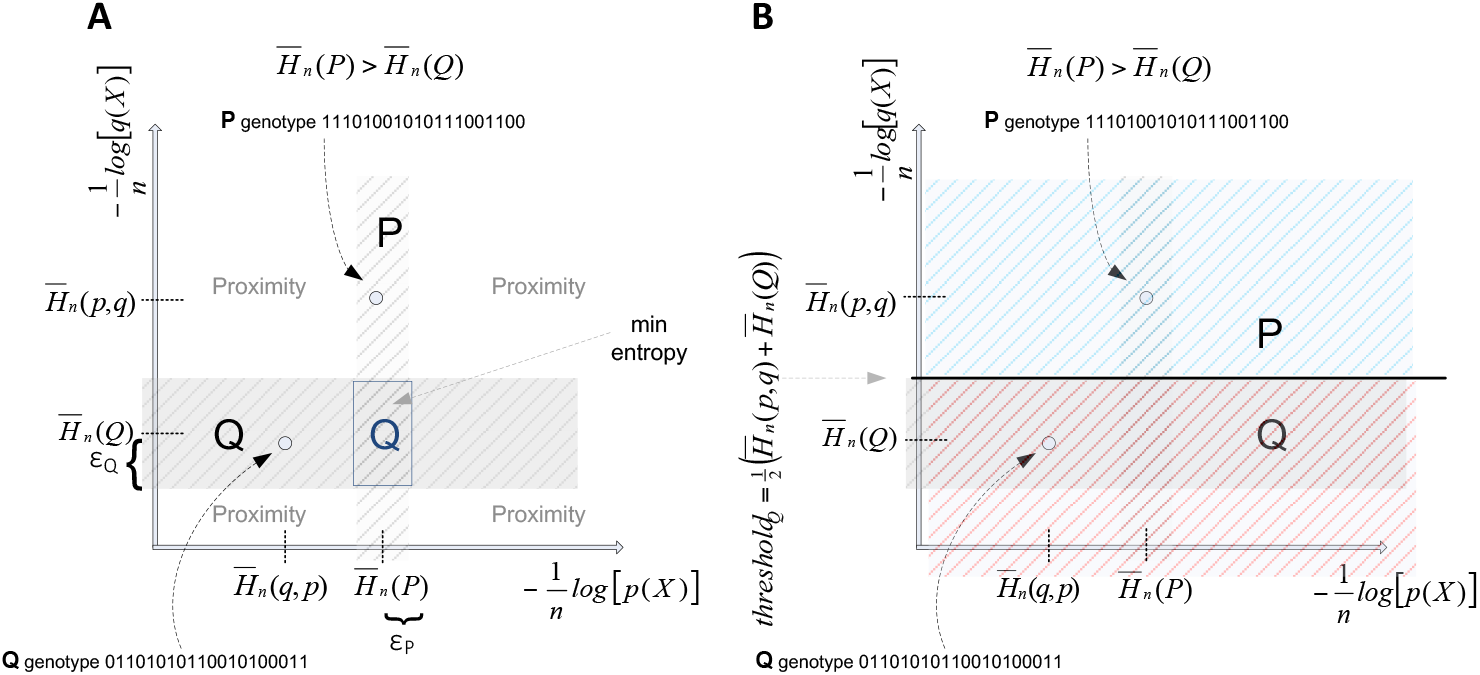
The naïve typicality classifier works according to exclusive typicality (with classification on min entropy in case of mutual typicality, and proximity to entropy rates in case of non-typicality). B: The simpler cross entropy classifier works by considering only the empirical entropy with respect to one population and classifying according to proximity to entropy rate vs. cross entropy rate.

Crucially, we show that given any arbitrary thresholds on SNP frequencies, the error rates are exponentially bounded and thus are asymptotically zero, as would be required from any classifier on high dimensional data, and the rate of decrease is proportional to the maximal of the two cross entropy criteria. A numerical simulation of the log-probability space and the resulting error rates in a scenario of differing population entropy rates is depicted in Fig. 13 (real worldwide distant populations often have different SNP-based diversities, as reflected by property ‘f’ in section *Properties of sequences of genetic variants*).

**Fig. 13:**
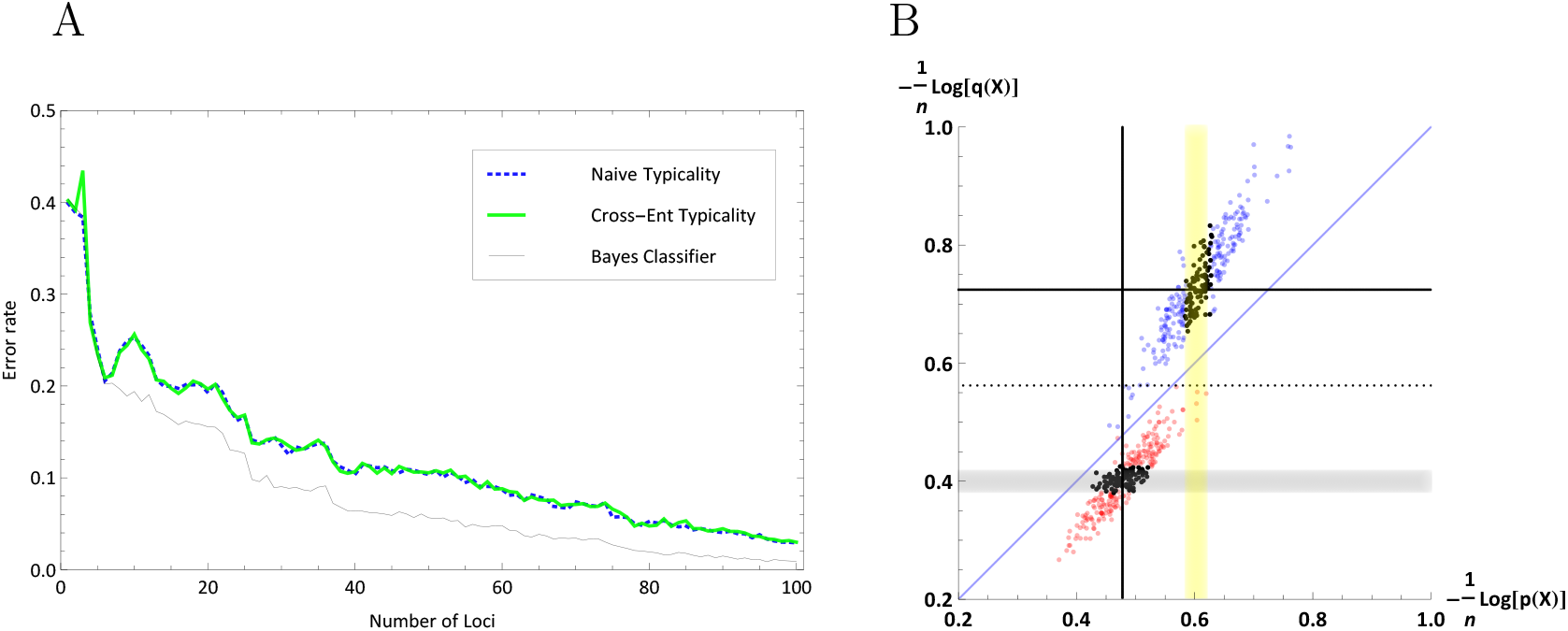
The performance of the typicality-based classifiers vs. an optimal Bayes classifier when population entropy rates differ (given known underlying allele frequencies). A: The error rates of the typicality classifiers demonstrate a good performance even for close populations. | B: The two clusters on the log-probability plot portray a strong horizontal separation (dotted line represents the cross entropy classification threshold), here at *n* = 300 SNPs (w/600 samples). In both panels SNP frequencies were modeled on Beta distributions (*α*_*P*_ = 4, *β*_*P*_ = 20, *α*_*Q*_ = 2, *β*_*Q*_ = 20) at each locus, with 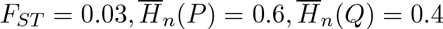.

Further simulations of the typicality classifiers reveal a low performance when the two cross entropy criteria are very similar (generally associated with similar population entropy rates, but not necessarily). A log-probability plot with respect to the cross entropy classifier reveals that this phenomenon is due to a relatively weak vertical/horizontal separation of the clusters (Fig. 14).

**Fig. 14:**
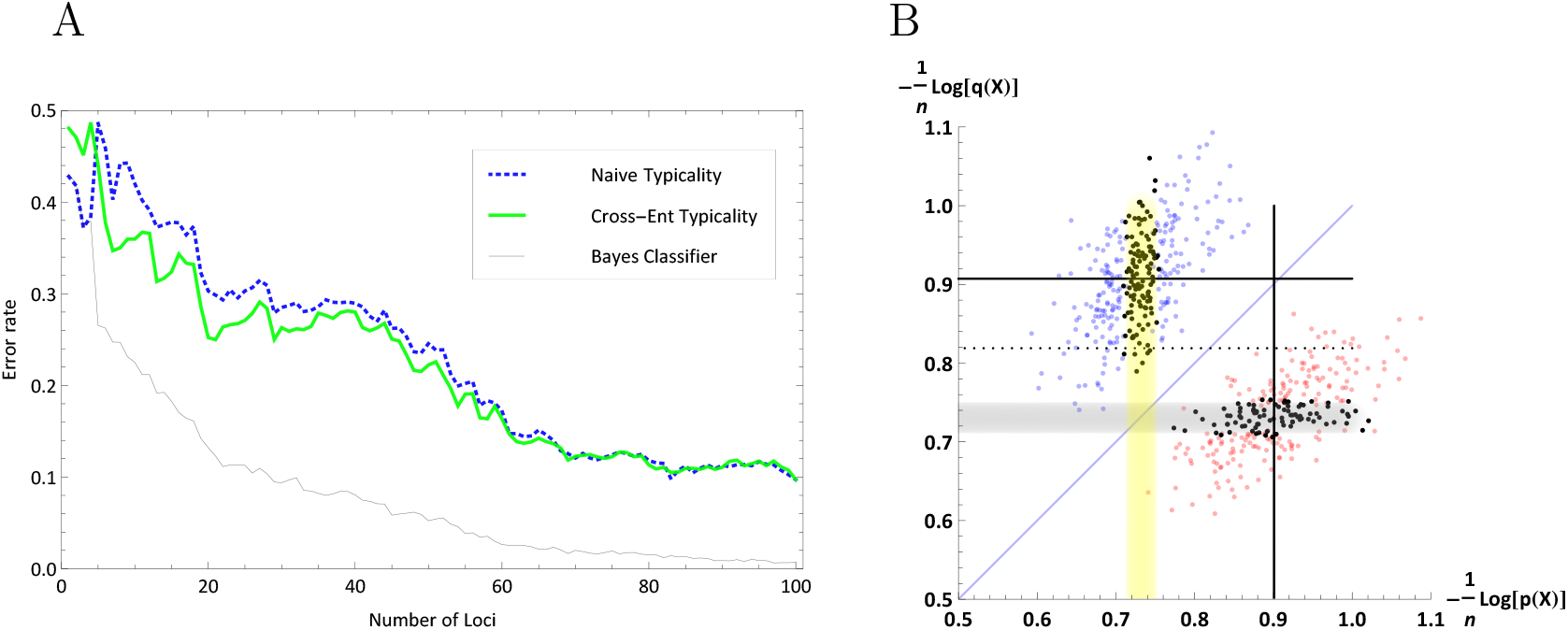
The performance of the typicality-based classifiers vs. an optimal Bayes classifier when population entropy rates are very *similar* (given known underlying allele frequencies). A: The error rates of the typicality classifiers demonstrate relatively poor performance. | B: The two clusters on the log-probability plot portray a weak horizontal separation (dotted line represents the cross entropy classification threshold) even at *n* = 200 SNPs (w/600 samples), while maintaining separation with respect to the Bayes classification line (thin blue). In both panels SNP frequencies were modeled on Beta distributions (*α*_*P*_ = 2, *β*_*P*_ = 6, *α*_*Q*_ = 2, *β*_*Q*_ = 6) at each locus, with 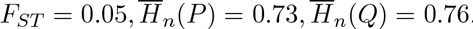.

### 4.4 Sampling Noise

The typicality classification models have been thus far defined parametrically, using the underlying frequencies of SNPs across the two populations. In practice, however, estimated frequencies from available data, rather than ‘true’ values must be used. This introduces a source of stochastic noise into our system. The link of noise to uncertainty was noted very early by [Shannon and Weaver, 1949], who stressed that: ‘If noise is introduced, then the received message contains certain distortions…[and] exhibits, because of the effects of the noise, an increased uncertainty.’ Fano’s Inequality provides a lower bound on the minimum error rate attainable by any classifier on symbols through a noisy channel, in terms of entropies and conditional entropies of the source and destination. Suppose that we know a random variable *Y* and we wish to guess the value of a correlated random variable *X*. We expect to be able to estimate *X* with a low probability of error only if the conditional entropy *H*(*X*|*Y*) is small. Assuming binary symbols as in our genetic framework, a simplified and slightly relaxed quantification of this idea is the lower bound on the error, *e* ≥ *H*(*X*) − *I*(*X*; *Y*)−1 ([Cover and Thomas, 2006]).

Simulations of a variety of classification methods on genetic data show that performance is degraded with smaller population samples, most notably for close populations ([Rosenberg, 2005]). Estimates of SNP frequencies computed at the training stage deviate from their true population values due to *statistical sampling*, a source of noise different from that introduced by error in the sequencing of ‘test samples’. This is the case even when genetic sequencing is 100% error free since it is purely a statistical effect.

Here we highlight a surprising feature of all typicality based classifiers under such training noise. For scenarios of close populations (low *F*_*ST*_), differing entropy rates and small training sample sizes, the typicality based classifiers consistently out-perform the Bayes classifier when allele frequencies are estimated using a natural (naïve) or maximum-likelihood estimator (MLE).^5^ Allele frequency estimates of zero are replaced with a small constant proportional with the sample size, a common procedure to avoid zero genotype frequencies ([Rosenberg, 2005]; [Phillips et al., 2007]). Specifically, for a sample of size *m*, the naïve ML estimator sets frequencies to be 1/(2*m* + 1) for counts of zero alleles, and 1 − 1/(2*m* + 1) for counts of *m* alleles (since we assume SNPs have some cut-off frequency), as in [Phillips et al., 2007]. The advantage of such an estimator is that it *makes no underlying assumptions* on the ‘true’ distributions of the parameters estimated (in particular, it makes no assumption on SNP frequencies being distributed i.i.d. across loci), i.e., no prior is utilized.^6^ We may also incorporate a Bayesian approach to allele frequency estimation by using a prior based on some justified model, effectively attenuating the sampling noise. A reasonable prior (close-to-optimal) can be produced by updating a histogram across a large number of loci, given the *assumption* of identically distributed frequencies across loci. In conjunction with the binomial likelihood function this results in a posterior distribution.^7^ These phenomena are illustrated in Fig. 15.

**Fig. 15:**
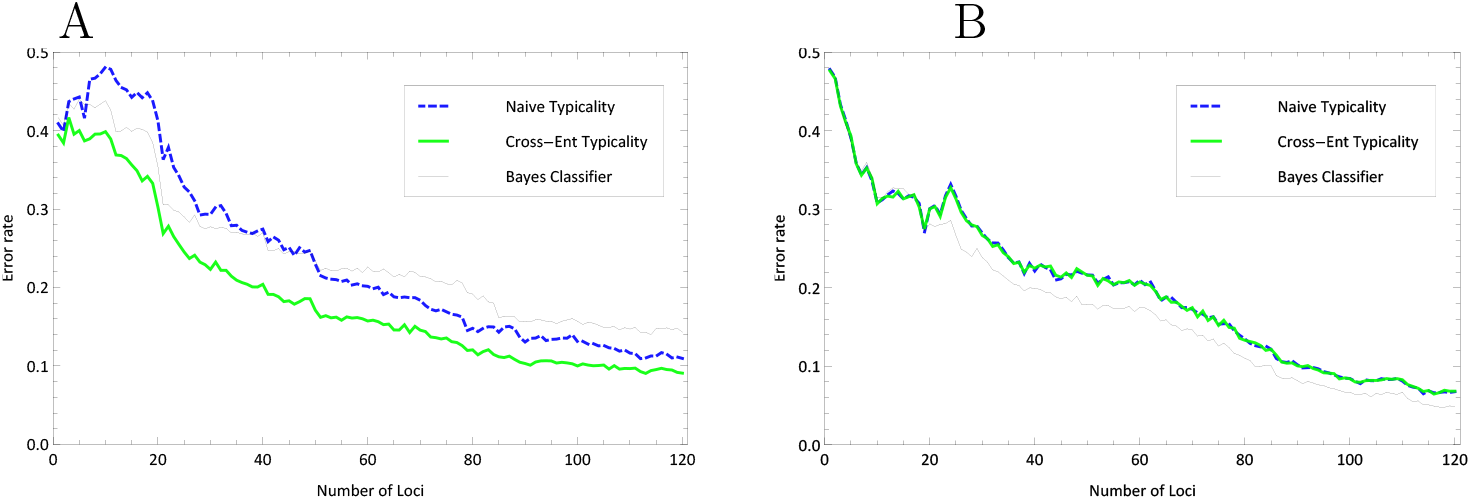
With maximum likelihood estimation of allele frequencies under small training sets (high sampling ‘noise’ level) and differing population entropy rates the typicality based classifiers consistently out-perform a Bayes classifier (Panel A), an advantage which dissipates if the ‘true’ prior is known and a Bayesian posterior is employed (Panel B). In both panels SNP frequencies were modeled on Beta distributions (*α*_*P*_ = 4/*β*_*P*_ = 20, *α*_*Q*_ = 2/*β*_*Q*_ = 20) at each locus, with 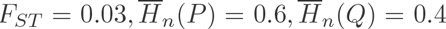, with a training set of 9 samples from each population, averaged over 6 training runs.

What is the underlying reason for the typicality classifiers’ resilience to training noise under a naïve maximum likelihood estimation of allele frequencies? From AEP considerations, the noisy samples from population *P* will cluster in the log-probability space around the coordinate 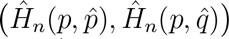, while the noisy samples from *Q* cluster around the coordinate 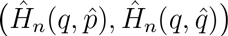, where 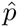 denotes the vector of length *n* such that 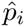 is the maximum-likelihood estimate of *p*_*i*_, and a similarly for 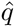. Simulations indicate that the introduction of sampling noise causes the population clusters to disperse, and more importantly, to shift towards the diagonal Bayesian separation line and therefore compromise the Bayes classifier’s accuracy (as can be appreciated from comparing the two panels of Fig. 16).

**Fig. 16:**
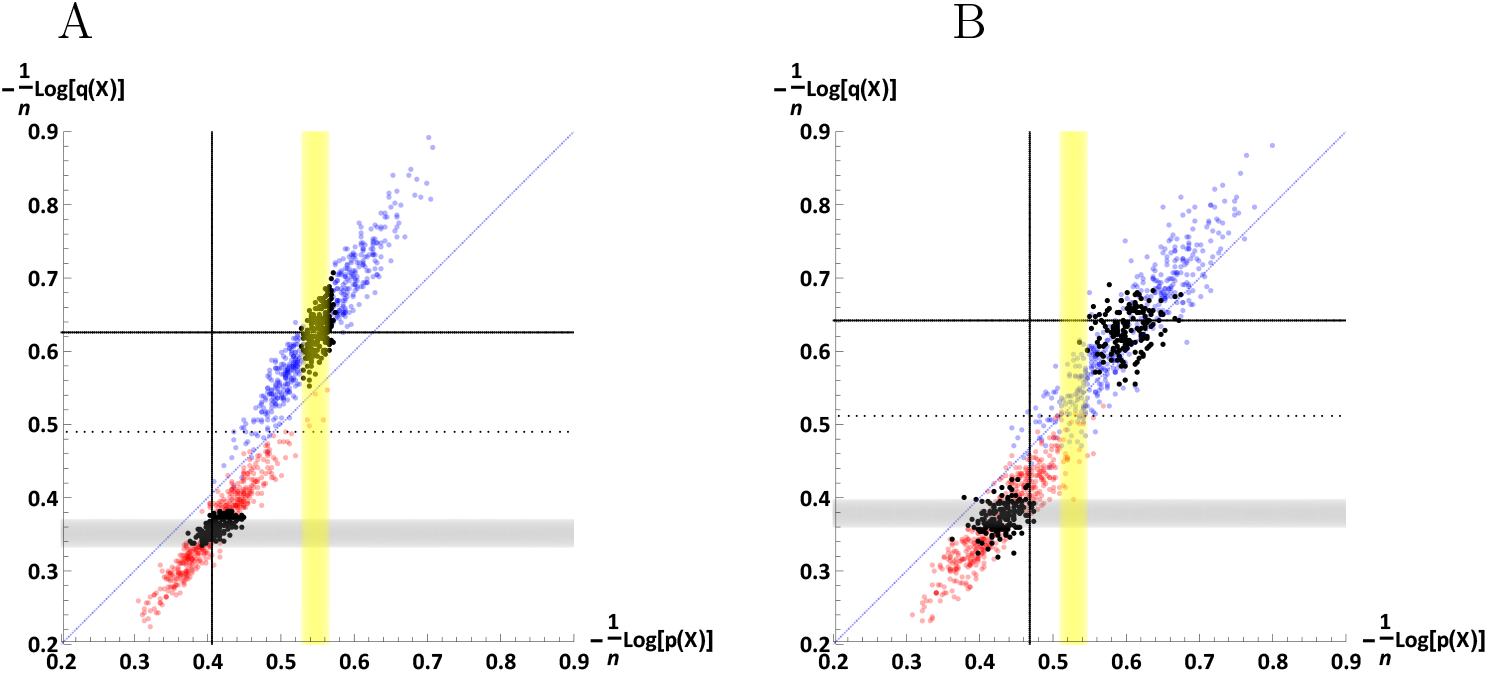
The effect of training noise on genotype samples on the log-probability plot. A: a scenario without sampling noise. | B: the same scenario when sampling noise is introduced (only 12 training samples from each population), resulting in better horizontal separation (cross entropy classifier) than a diagonal one (Bayes classifier). 1200 samples were drawn from each population at *n* = 300 SNPs, where population SNP frequencies were modeled on Beta distributions for *P* and *Q* with *α*_*P*_ = 6/*β*_*P*_ = 40, *α*_*Q*_ = 3/*β*_*Q*_ = 40, at each locus.

Formally, from Jensen’s inequality we get,

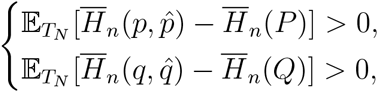

where 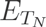 denotes the expectation value with regard to a training scenario of sample size *N*.

We now turn to the resilience of the typicality classifiers and consider the effect of noise on the cross entropy classifier, where without loss of generality, *C*_*Q*_ > *C*_*P*_. Note that,

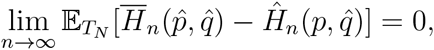

since 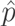 is an unbiased estimator of *p*. Note that because both *p*_*i*_ and *q*_*i*_ are distributed i.i.d., it holds for all *i*: 1,…,*n* that

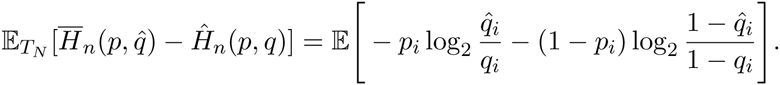

Heuristically, this difference is likely to be much larger than the difference

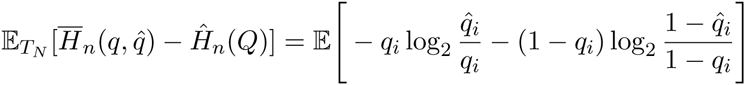

for the following reason: in both cases a large contribution to the difference comes from where *q*_*i*_ is small and 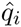 provides an underestimate for *q*_*i*_, resulting in a large logarithm 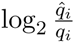. However, in the second difference, this logarithm has a prefactor *q*_*i*_ which is small, whereas in the first difference the prefactor *p*_*i*_ which on average is significantly larger.

A similar type of argument suggests that the difference 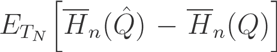 is relatively small compared to 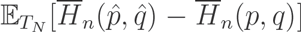. These heuristics make plausible that the threshold of the cross entropy classifier, calculated as the average of 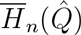 and 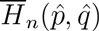, still separates well the ‘noisy’ clusters, for which the vertical coordinates are given by 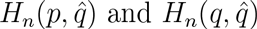.

### 4.5 Relative-entropy typicality

A well-known extension of the concept of typical-set is the ‘relative entropy typical set’ ([Cover and Thomas, 2006], Section 11.8). For any fixed *n* and *ε* > 0, and two distributions *P*_1_ and *P*_2_, the relative entropy typical set 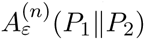 entails all sequences of length *n* such that,

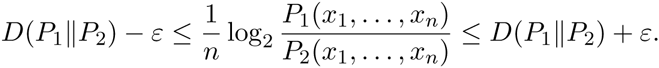

Similar to standard set typicality, the relative entropy typical set asymptotically includes all the probability,

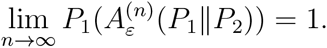

Crucially for our purposes there exists an associated AEP theorem for relative typicality ([Cover and Thomas, 2006], Theorem 11.8.1): Let *X*_1_, *X*_2_,…,*X*_*n*_ be a sequence of random variables drawn i.i.d. according to *P*_1_(*x*) and let *P*_2_(*x*) be any other distribution on the same support, then,

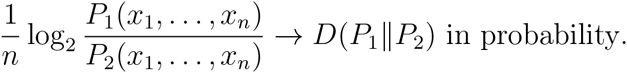

However, to account for the non-stationary sources (i.e. the variation of SNP frequencies across loci, a standard feature of population data), as in our treatment of entropy typicality, we need to modify the definition of relative-entropy typicality and derive an associated AEP theorem (Appendix B.4).

We may now construct a naïve classifier based on exclusive relative-typicality, with some choice of an epsilon margin around the respective KL-Divergence rate, and some means of resolution for the cases of mutual relative-typicality or lack of relative-typicality. Alternatively, a more straightforward construction is to simply to classify by proximity to the respective KL-Divergences,

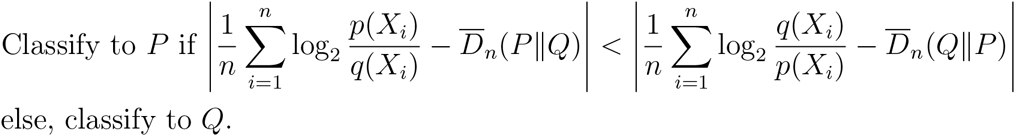

Where the KL-Divergence rate is defined in Eq. (13). Fig. 17 is a schematic of such classifiers with respect to the log-probability space. (see Appendix A.4 for a closed-form formulation of the error rate).

**Fig. 17:**
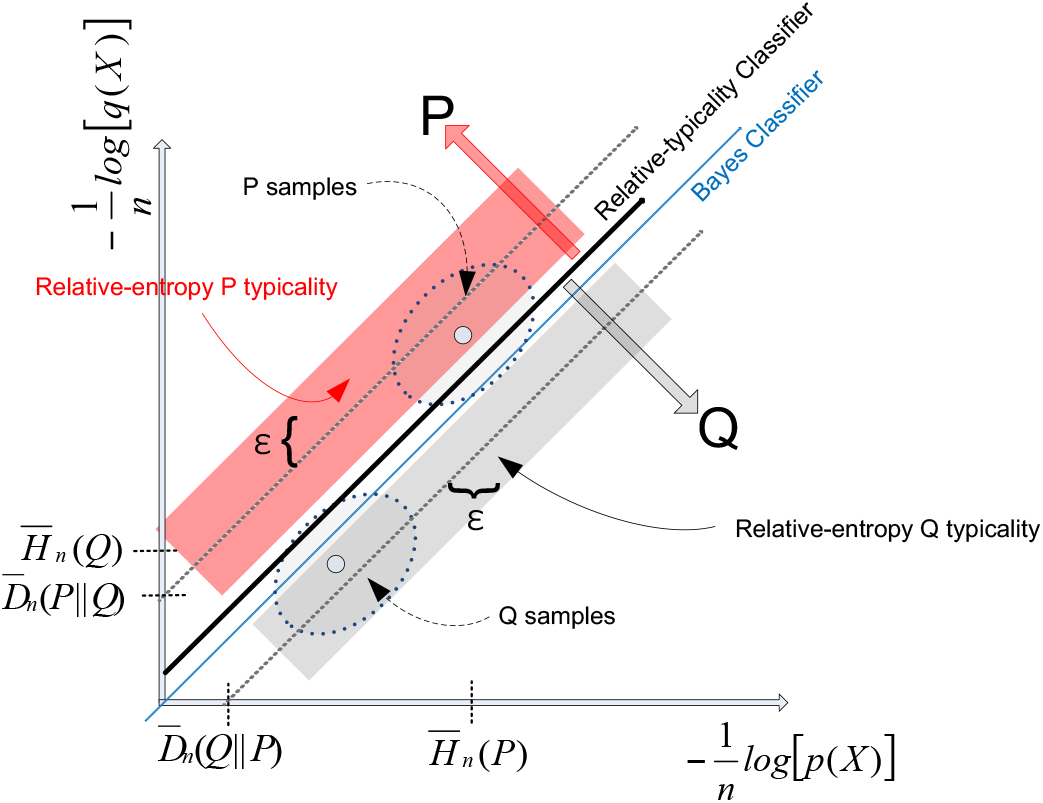
A schematic representation of a straightforward implementation of a proximity-based relative entropy typicality classifier (black diagonal line) and a naïve relative-entropy classifier (dotted diagonal lines), with respect to some arbitrary epsilon (dark stripe margins, red for *P* and grey for *Q*). The proximity-based relative entropy classifier merges in performance with a Bayes classifier with an uninformative class prior (blue) line only when 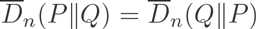, and is represented by the line 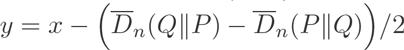.

Finally, note that this classifier can also be described as,

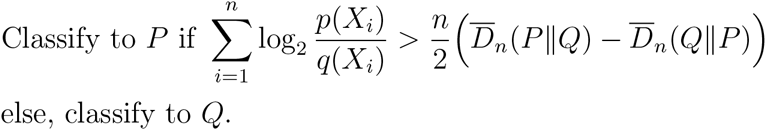

While on the other hand, a Bayes classifier with prior *α* classifies as follows,

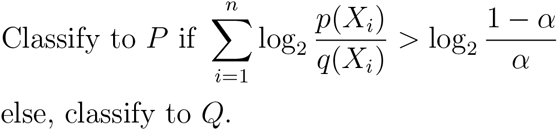

Hence, the relative entropy classifier that classifies by proximity, as described above, is exactly a Bayes classifier with prior *α*, where *α* satisfies,

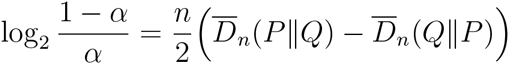

that is,

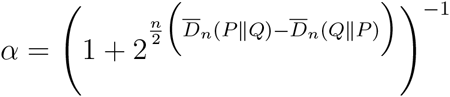

where different choices of ‘*ε*’ would correspond to choosing different priors for the Bayes classifier. Not surprisingly, the relative-entropy classifier is similarly not resilient to learning-based noise.

## 5 Discussion

> Simplicity is the final achievement.
>
> — -- F. Chopin.

The availability of high-throughput SNP genotyping and the nature of polymorphisms across loci and diverse populations suggest a fruitful application of one of the core ideas in information theory, that of set-typicality and its associated properties. In this treatment, we have employed conceptual and formal arguments along with evidence from numerical simulations to demonstrate that long sequences of genotype samples reveal properties that are strongly suggestive of *typical sequences*. This allowed us to produce versions of the asymptotic equipartition property that comply with population genetic data and consequently define the notion of mutual typicality and describe information-theoretic classification schemes. We do not claim here priority in invoking the concept of typical sets broadly in biology. In examining the fitness value of information, [Donaldson-Matasci et al., 2010] have made use of the asymptotic properties of typical sequences to capture properties of typical *temporal* sequences of selection environments and their payoffs in evolution. However, our use of a typical-set framework to analyze long *sequences of genetic variants* is, as far as we know, original. Moreover, to the best of our knowledge, a general analysis of mutual and exclusive typicality and the interplay of multiple typical sets (from sources defined on the same space) is another original contribution. In this context, we note that the related notion of ‘strong typicality’ is only applicable for stationary sources where the sample frequency of a symbol is closely linked to its underlying distribution, and therefore not directly applicable in our framework, where alleles are not identically distributed across loci.

The consideration of noise as a source of classification error, and a subsequent quantification, is of course, not new. From a machine learning perspective, one of the early insights of information theory was to consider a classification problem as a noisy channel. Fano’s inequality provides a lower bound on the minimum error rate attainable by any classifier on symbols through a noisy channel, in terms of entropies and conditional entropies of the source and destination. Suppose that we know a random variable *Y* and we wish to guess the value of a correlated random variable *X*. We expect to be able to estimate *X* with a low probability of error only if the conditional entropy *H*(*X*|*Y*) is small. Assuming binary symbols as in our genetic framework, a simplified and slightly relaxed quantification of this idea is the lower bound on the error ([Cover and Thomas, 2006]), 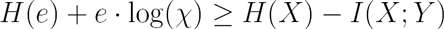.

Shannon (1956) has famously cautioned against jumping on ‘the bandwagon’ of information theory whose basic results were ‘aimed in very specific direction…that is not necessarily relevant to such fields as psychology, economics, and other social sciences’. He stressed that while ‘Applications [of information theory] are being made to biology…, A thorough understanding of the mathematical foundation and of its communication application is surely a prerequisite to other applications…’, finally concluding that, ‘I personally believe that many of the concepts of information theory will prove useful in these other fields – and, indeed, some results are already quite promising – but the establishing of such applications is not a trivial matter of translating words to a new domain, but rather the slow tedious process of hypothesis and experimental verification.’

Notwithstanding Shannon’s concerns, there have been numerous attempts at borrowing both informational concepts and technical results from information theory in the biosciences. In a recent illuminating review, [Vinga, 2014] highlights several information-theoretic measures that have been applied widely, e.g., to compare sequences in an alignment-free context, provide block-entropy and complexity estimation, and assess DNA sequence compression limits. [Ulanowicz et al., 2009] has ushered in the “return of information theory” by using conditional entropy to quantify sustainability and biodiversity. [McCowan et al., 2002] had emphasized the prominent role of noise in “constraining the amount of information exchanged between signallers and perceivers” in ecological and social contexts and for signal design and use. By applying quantitative and comparative information-theoretic measures on animal communication, they hoped to provide insights into the organization and function of “signal repertoires”. Similarly, [Levchenko and Nemenman, 2014] have shown how cellular noise could be quantified using mutual information, and the implications of measuring such noise in bits. Even more recently, [Lan and Tu, 2016] have focused on the “inherent noise in biological systems’ which they have argued can be analyzed by ‘using powerful tools and concepts from information theory such as mutual information, channel capacity, and the maximum entropy hypothesis”, with subsequent analysis mostly restricted to entropy and mutual information in their capacity as statistical measures. Other authors have made claims, admittedly of a conjectural nature, on the relevancy of information theoretic results to principles of evolution and genetic inheritance. For instance, [Battail, 2013] has claimed that the trend of biological evolution towards increasing complexity and hereditary principles requires the implementation of error correcting information-theoretic codes, which are inevitable and ‘logically necessary’ once it is clear that ‘heredity is a communication process’, while at the same time emphasizing that these are ‘merely speculations’.

While some of these and other approaches have been interesting and insightful, the conceptual and formal link to information theory mainly comprises of metaphoric use of otherwise technical information theoretic concepts and terms, such as communication channel and noise, or the employment of quantitative measures of variation and dependency that originate in information theory. Indeed, in a review of the contribution of information theory to molecular biology, [Fabris, 2009] concludes that the evidence indicates the contribution is “no more than [on] a purely syntactic level” and wherever use of a statistical framework is required, then “tools such as mutual information, entropy and informational divergence, can be used with profit”. The author further conjectures that this is due to a naive “assumption of a substantial equivalence between the Shannon unidirectional transmission system and the DNA-to-protein communication system.”

### 5.1 Channel capacity

The concept of *channel capacity*, which also plays a central role in communication theory, may serve to further highlight the shared properties identified here between long sequences of symbols generated by a random source and communicated across a noisy channel, and long genotypes originating from a natural population. The channel capacity is the tight upper bound on the rate at which information can be reliably transmitted over a noisy communications channel. The usefulness of this notion in other domains was famously identified by [Kelly, 1956]. Kelly analyzed a scenario which seems to possess the essential features of a communication problem: a gambler that utilizes the received symbols of a noisy communication channel in order to make profitable bets on the transmitted symbols. Kelly then demonstrated that, just as information can be transmitted over a noisy communication channel at or near Shannon’s channel capacity with negligible error, so can this gambler compound his net worth at a maximum rate with virtually no risk of ‘total loss’, equal to the mutual information of the source and receiver (by apportioning his betting amount precisely according to the noise level for each symbol).

More formally, the “information” channel capacity *C* of a discrete memoryless channel with respect to sources *X* with alphabets supported on *χ* and consequent outputs *Y* with alphabets supported on *y*, is an inherent property of the channel such that, *C* = max_*P*(*X*)_ *I*(*X;Y*), or for nonstationary sources representing our population model,

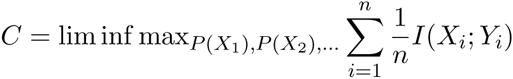

where the maximum is taken over all possible distributions *P*(*X*_*i*_) of the source ([Verdu and Han, 1994]). The capacity is commonly interpreted as the highest rate in bits per channel use at which information can be sent with arbitrarily low probability of error. Shannon’s channel coding theorem then relates the maximum information rate possible across the channel with its capacity ([Cover and Thomas, 2006], Ch.7).

We now propose an analogy between the effect of communication noise on channel capacity and the effect of sampling noise on classification accuracy, centered on around the mutual information between inputs and outputs. If we interpret *X* as a random variable representing the *n*-SNP genotype from the pooled source populations and *Y* as a random variable representing its source population, then the mutual information *I*(*X; Y*) captures the *informativeness* of the set of *n* markers for population assignment (see [Tal, 2012a], [Tal, 2012b] for the multilocus formulation). This is also known as the *Infomax principle* in feature selection, where a subset of features is chosen so that the mutual information of the class label given the feature vector is maximized ([Rosenberg et al., 2003]; [Zhao et al., 2013]; see [Peng et al., 2005] for the *Max-Dependency* principle). If we now take the *informativeness I*(*X; Y*) to represent the *maximal information extractable* across all possible classifiers, a workable analogy with communication-based channel capacity, which is also expressed in terms of mutual information, becomes evident. Under this interpretation, the *inferential channel capacity* is achievable by the optimal Bayes classifier, under known distribution parameters ([Hastie et al., 2009]), i.e., in the absence of sampling noise; otherwise, given any finite sample size at the learning stage, there may be no single classification scheme that obtains maximal performance under all data scenarios. Indeed, the lack of a universally best model for classification is sometimes called the *no free lunch theorem*, which broadly implies that one needs to develop different types of models for different data, since a set of assumptions that works well in one domain may work poorly in another ([Murphy, 2012]).

### 5.2 Dimensionality reduction

It is worthwhile highlighting an additional feature of our log-probability space, with possible pragmatic use. The mapping of genotype samples to the log-probability space shares some core features with standard dimensionality reduction schemes such as PCA, which are often deployed for visualization purposes or as pre-processing in the context of unsupervised learning. Most prominently, [a] the effect of higher dimensionality (larger *n*) on cluster separability, [b] the effect of population differentiation (*F*_*ST*_) on cluster proximity, [c] the effect of distribution entropy rates on the cluster shape, and [d] the general effect of a possible presence of LD given the explicit (implicit, in the case of PCA) assumption of LE. At the same time, the log-probability perspective provides information with respect to a supervised learning framework, most prominently by revealing the effect of noise in the training stage on the clusters of ‘test samples’, and on the estimated quantities employed by an information-theoretic oriented classifier, such as our cross entropy typicality classifier.

### 5.3 Linkage Disequilibrium

When populations have some internal structure (deviation from panmixia) then loci are in linkage disequilibrium (LD). In terms of the communication framework, LD is analogous to time-dependency of symbols generated by the source, such that the channel is no longer *memoryless*. How will our results fare when such dependencies are introduced into the inferential framework?

Previous work on analogies and implementations of information theoretic concepts has highlighted this difficulty. For instance, in his famous approach to betting strategies from an information-rate perspective, [Kelly, 1956] has also emphasized that in the presence of time-dependency of symbols the results he obtained may no longer be relevant, acknowledging that ‘theorems remain to be proved’ if the symbol transmission entails dependency on time or past events.

Since our results are intrinsically based on AEP theorems, we would be interested to pursue some generalization of the AEP for (nonstationary) sources with *dependent* symbols. The Shannon-McMillan-Breiman theorem ([Cover and Thomas, 2006]) is an extension of Shannon’s original AEP for discrete i.i.d. sources, and holds for discrete-time finite-valued *stationary ergodic sources*, which in general have dependent symbols. However, the closest to general nonstationary sources with dependent symbols for which an AEP holds are a class of nonstationary sources called ‘asymptotically mean stationary’ or AMS sources ([Gray, 2011]). These are sources which might not be stationary, but are related to stationary sources in a specific way. Such sources are equivalent to sources for which relative frequencies converge and bounded sample averages converge with probability one, but not necessarily to simple expectations with respect to the source distributions. They include, for example, sources with initial conditions that die out (asymptotically stationary sources) along with sources that are block-stationary, e.g., extensions of the source are stationary.

Crucially for our purposes, general patterns of LD found in population SNP data should not be expected to conform to the specific properties characteristic of AMS sources, and therefore we cannot expect an AEP to hold for such data. Nevertheless, we would like to see whether a ‘naïve’ approach to classification by typicality, akin to that taken by the naïve Bayes, might still be productive. Adopting such ‘naïve’ approach means that we employ the same expressions for genotype probabilities, empirical entropies, population entropy and cross entropy rates, which had all assumed statistical independence.^8^

Numerical analysis shows that with various patterns of LD the typicality classifiers do not account well for its presence, contrary to the naïve Bayes classifier. Under any type of LD, clusters on the 2D log-probability plot tend to substantially disperse (elongating diagonally), breaching the typicality threshold even for very large *n* where we would expect substantial separation (Fig. 18). Interestingly, this diagonal elongation gives a new perspective on the well-known phenomenon by which under LD naïve Bayes classifiers still outperform far more sophisticated alternatives, and make it surprisingly useful in practice even in the face of such dependencies ([Hastie et al., 2009] section 6.6.3). We stress here that the increased dispersion of samples when LD is introduced *cannot* be taken as indicative of the well-known result that there is no AEP for nonstationary sources with dependent symbols, since samples are mapped to this space according to ‘naïve’ independence assumptions. Estimating the actual genotype probabilities (along with *joint* entropies and cross-entropies under these assumptions for constructing the typicality classifier) is beyond the scope of the models used in these simulations.

**Fig. 18:**
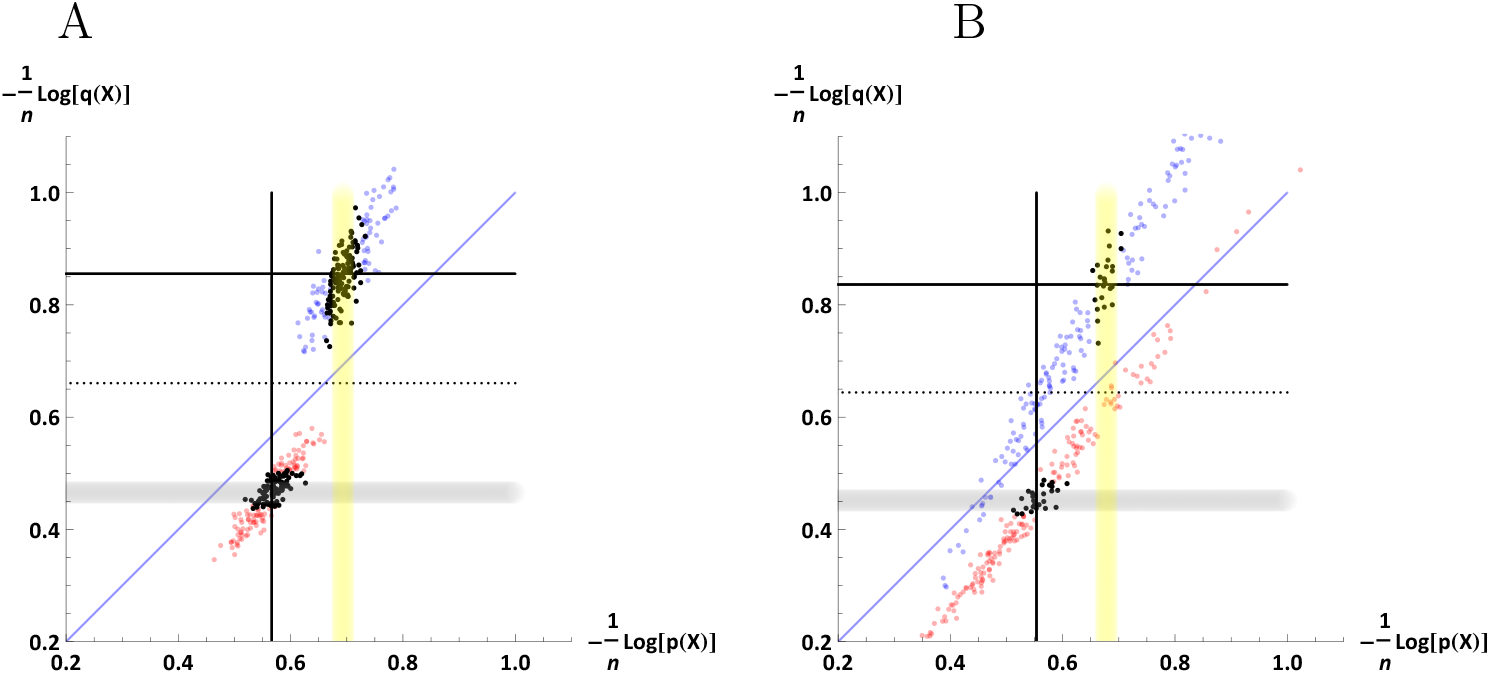
Clusters of genotype samples from the two populations are elongated diagonally as a function of the amount of LD and its nature, substantially breaching the typicality classification threshold (dotted line) while maintaining separation with respect to the Bayes classification line (thin blue). Here 400 samples were drawn from two populations modeled under Beta distributions with *n* = 600 SNPs, *F*_*ST*_ = 0.04, with differing population entropy rates, with *ε* = 0.02 for typicality. A: No LD. | B: Moderate levels of LD.

## 6 Conclusion

There has recently been revived interest in employing various aspects of information theory for characterizing manifestations of information in biology. Arguably, quantitative analysis of biological information has thus far only superficially drawn from the groundbreaking ideas and formal results of this highly influential theory. Here, we have ventured beyond the mere utilization of information-theoretic measures such as entropy or mutual information, to demonstrate deep links between a core notion of information theory, along with its properties and related theorems, and intrinsic features of population genetic data. We have demonstrated that genotypes consisting of long stretches of variants sampled from different populations may be captured as *typical sequences* of nonstationary symbol sources that have distributions associated with population properties. This perspective has enabled us to treat typical genotypes as proxies for diverse source populations, analyse their properties in high dimensions and consequently develop an information theoretic application for the problem of ancestry inference. We hope that this work will open the door for further inquiry into the prospects of rigorous implementation of both ideas and technical results from information theory in the field of population genetics and biology in general.

The Mathematica code for generating the numerical simulations for the figures can be made available by request from the corresponding author.

## Acknowledgements

We would like to thank Jürgen Jost for his interest and constructive feedback on these ideas. We appreciate the input of Robert M. Gray on AMS sources. Special thanks also to Slava Matveev, Guido Montúfar and Michael Lachmann for some fruitful technical discussions. Finally, we acknowledge the Max Planck Institute for Mathematics in the Sciences for the platform to present these ideas in an internal seminar and for its generous support.

## A Appendix A

### A.1 Closed-form formulation of the naïve typicality classifier error rate

The error rate of the naive typicality classifier can be expressed as,

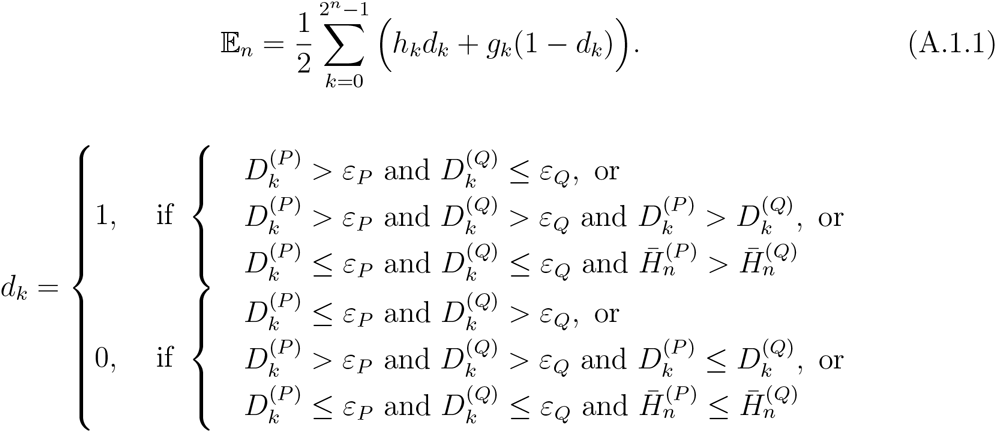

where

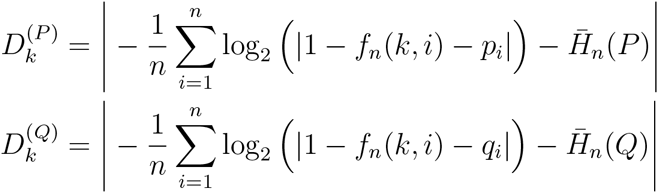

and where the genotype probabilities *h*_*k*_ and *g*_*k*_ and the indicator function *f*_*n*_ are defined as in ([Tal, 2012b], section 3.2),

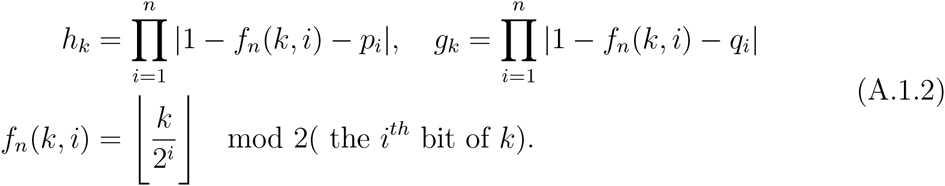

### A.2 Closed-form formulation of the cross entropy classifier error rate

The error rate of the cross entropy typicality classifier can be expressed using 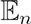 of Eq. (A.1.1) in conjunction with,

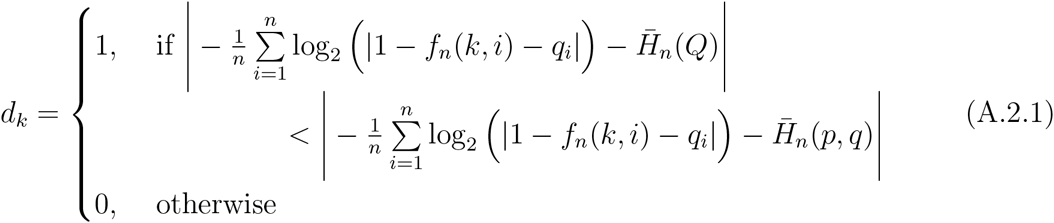

for the case where *C*_*Q*_ > *C*_*P*_, and similarly expressed in terms of the parameters of *P* when *C*_*Q*_ ≤ *C*_*P*_.

### A.3 Closed-form formulation of the generalization error of the cross entropy classifier under MLE

The expected test error *E*_*n,m*_ under all training samples of size *m* = {*m*1, *m*2} is an expectation over the conditional (on a particular sample of size m) test error 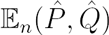 ^9^,

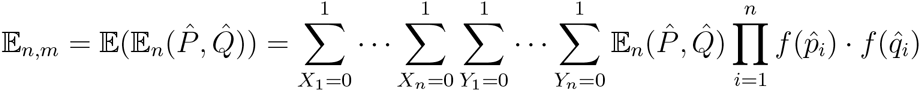

where we denote 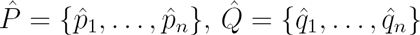.

Following the formulation in Eq. (A.1.1) we have,

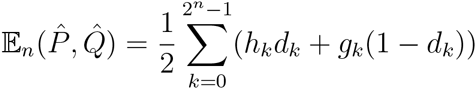

where the cross entropy classifier of Eq. (A.2.1) (for the case *C*_*Q*_ > *C*_*P*_) is expressed as conditional on a particular sample,

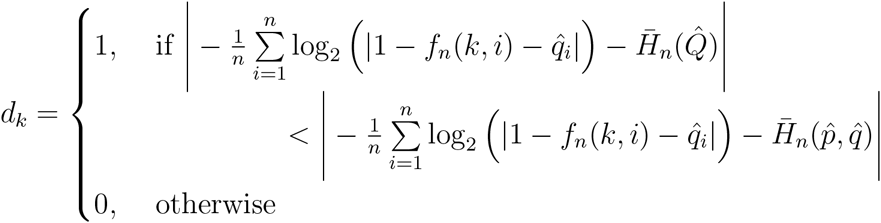

where 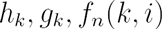 are defined with respect to the true frequencies, as in Eq. (A.1.2).

### A.4 Closed-form formulation of the error rate of the relative-entropy classifier

Following the formulation in Eq. (A.1.1) the error rate of the relative-entropy classifier can be expressed as,

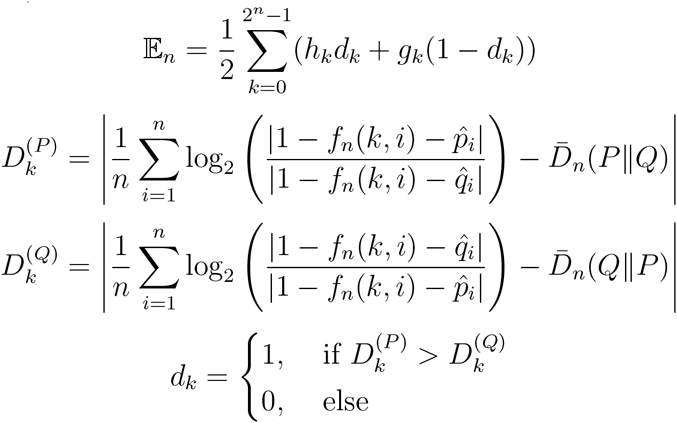

where the genotype probabilities *h*_*k*_ and *g*_*k*_ and the indicator function *f*_*n*_(*k*, *i*) are as defined in Eq. (A.1.2).

Note that the counterpart classifier-expressions for a Bayes (or maximum likelihood) classifier would in a corresponding formulation be expressed as a simple comparison of genotype probabilities,

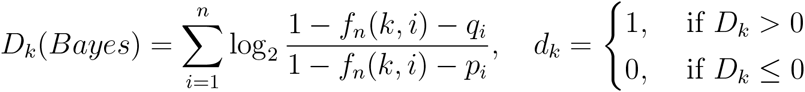

## B Appendix B

### B.1 Entropy and cross entropy rates

In this section we consider the expectation of entropy and cross entropy rates and their properties.

First, we recall some properties of a Beta distribution. Let *Y* ~ *B*(*α, β*). Then

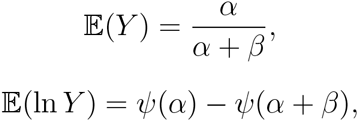

where 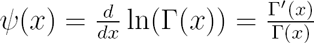 is a digamma function. Moreover, we have

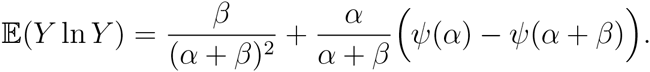

In fact, note that *Y* ~ *B*(*α, β*) implies that 1 ‒ *Y* ~ *B*(*β, α*). Therefore

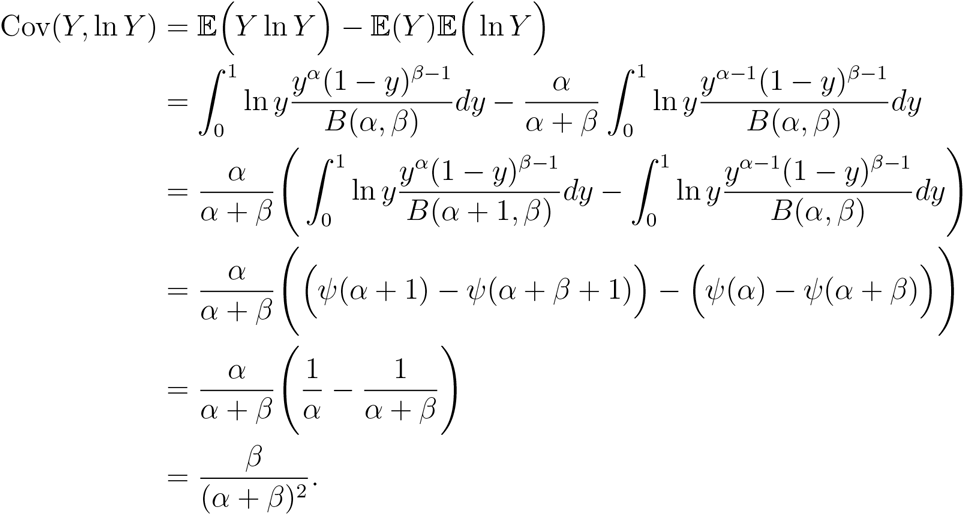

Therefore

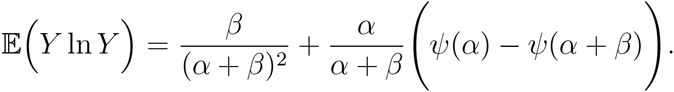

Suppose *p*_*i*_ and *q*_*i*_ are distributed i.i.d. according to 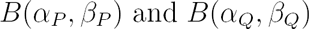 respectively. Then

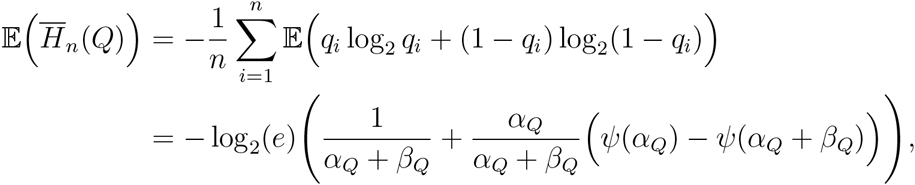

and similarly,

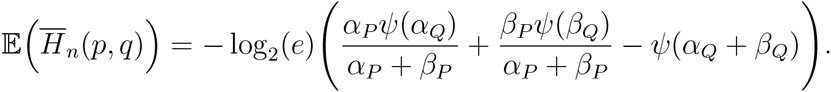

### B.2 cross entropy criteria

In this section of Appendix, we consider the cross entropy criteria 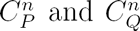 and its asymptotic properties. First, we have

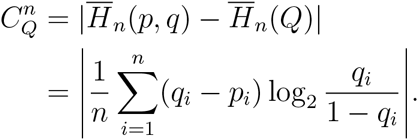

Assume that *p*_*i*_, *i* = 1,2,…, sampled by a random variable *X* with distribution 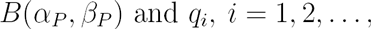 sampled by another independent random variable *Y* with distribution 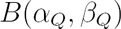. Then, by the law of large number we have the asymptotic property

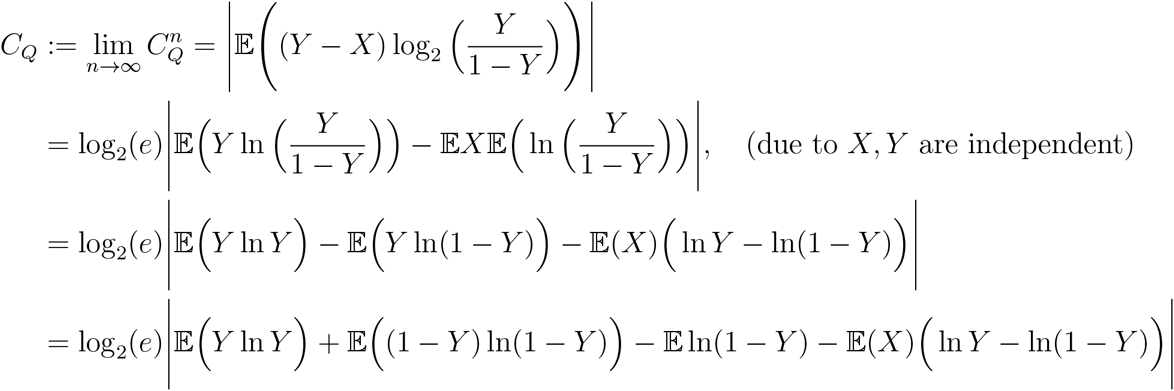

It implies that

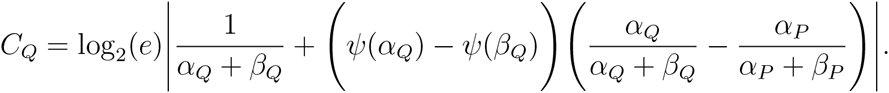

And similarly we also obtain

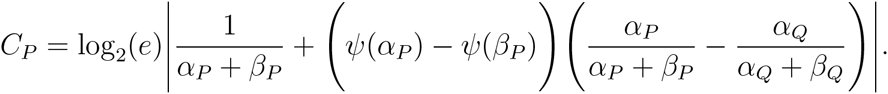

Then we have immediately some corollaries:

**Corollary B.2.1.** *C*_*Q*_ = 0 *if and only if*

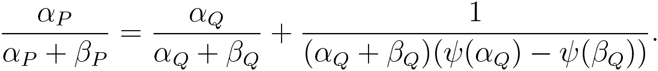

*Note that this equation has a lot of solutions* (*e.g. α*_*P*_ = 2, *β*_*P*_ = 10, *α*_Q_ = 2, *β*_*Q*_ = 4).

**Corollary B.2.2.** 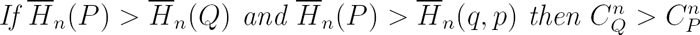.

*Proof*. In fact, we have

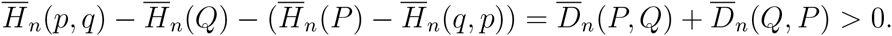

It implies that

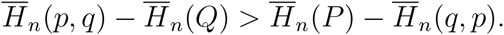

Moreover, due to the second condition, we have 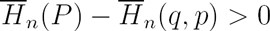. Therefore,

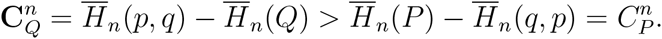

It completes the proof.

**Corollary B.2.3.** *Assume that* 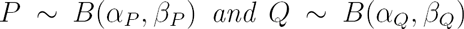 *satisfying* 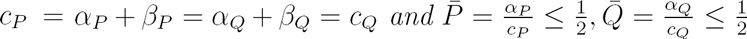. *If furthermore* 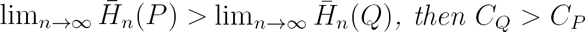.

*Proof*. In fact, it is enough to prove that for large enough *n* we have 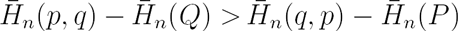. Indeed, note that

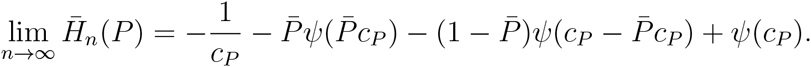

Therefore, the condition 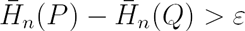 for all *n* implies that

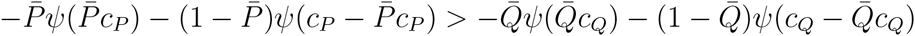

which implies that 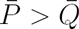.

Also we have

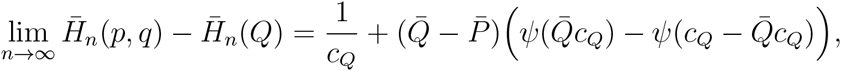

and 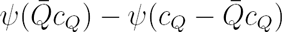 is decreasing with respect to 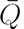. It implies the proof.

**Remark B.2.1.** *If C*_*P*_ = *C*_*Q*_ = 0 *then*

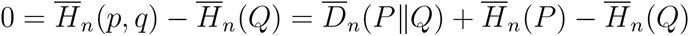

*and*

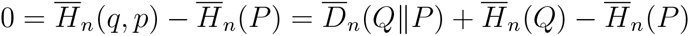

*This implies that 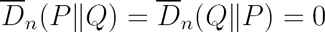 which happens if and only if P = Q.*

### B.3 Normalized pairwise distances

In this section, we first consider the average normalized pairwise distance in the set of all sampled genotypes and in the set of typical ones. We consider both the stationary and the non-stationary case.

#### B.3.1 Stationary case

**In** the stationary case *p*_*i*_ = *p* for all *i* = 1,…, *n* we have some first geometric properties of typical set as follows. Given *ε* > 0 and 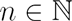, denote by

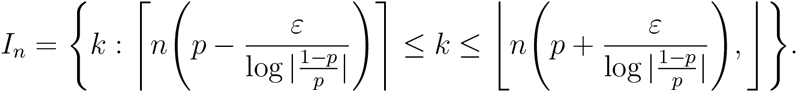

Then

(i)

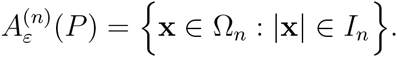

(ii)

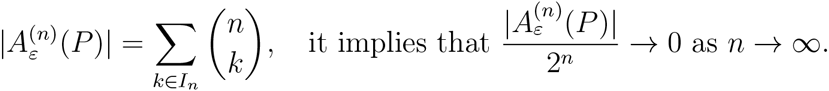

(iii)

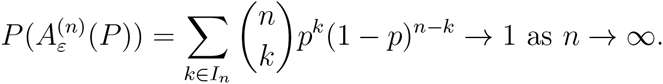

(iv)

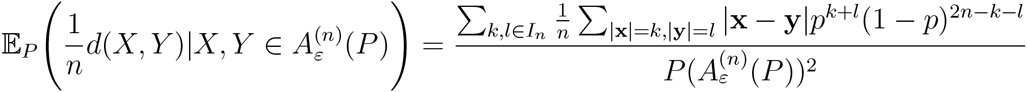

(v)

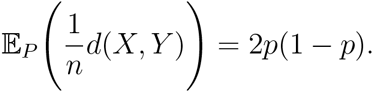

Let *C* be the centroid of Ω_*n*_ corresponding to distribution *P*, i.e. *c*_*i*_ = *p*_*i*_ for all *i* = 1,…, *n*. We also have a nice following property

**Proposition B.3.1.** *The covariance between the normalized generalized Hamming 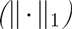 distance between X and C with respect to the Euclidean distance of their corresponding points in log-probability coordinate is non-negative, i.e.*

*(a)*

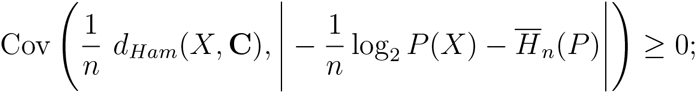

*(b) Equality holds if and only if 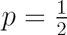;*

*(c) as n goes to infinity, this covariance goes to zero;*

*(d) when the entropy rate increases, the covariance decreases;*

*(e) statements in (a)-(d) are also true for correlation.*

*Proof*. First of all, note that in this case

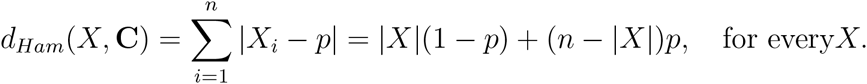

Therefore, it is easy to obtain

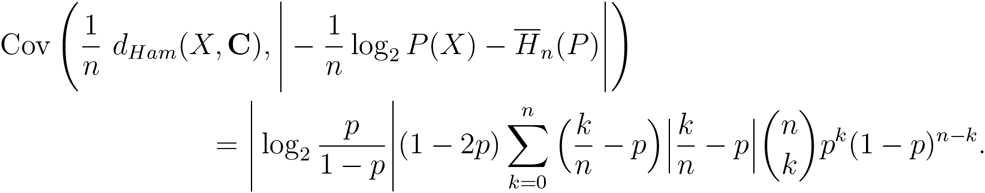

Put

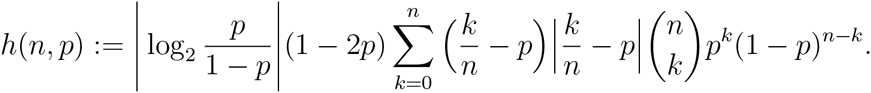

It is also easy to see that *h*(*n,p*) = *h*(*n*, 1 – *p)*. Without loss of generality, we assume that 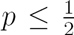. When 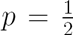, the covariance is zero. Moreover, we can prove that *h*(*n,p*) decreases in 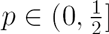 and in *n.*

It implies the proof.

#### B.3.2 Non-stationary case

Now we consider the non-stationary case. First, denote by *D*_*n*_(*X, Y*) the normalized Hamming distance of two genotypes *X* and *Y*, i.e.

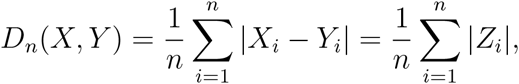

where *Z*_*i*_ is a random variable which is 1 with probability 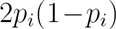 and 0 with probability 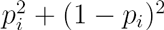.

Then the expectation and variance of *D*_*n*_ can be easily calculated as

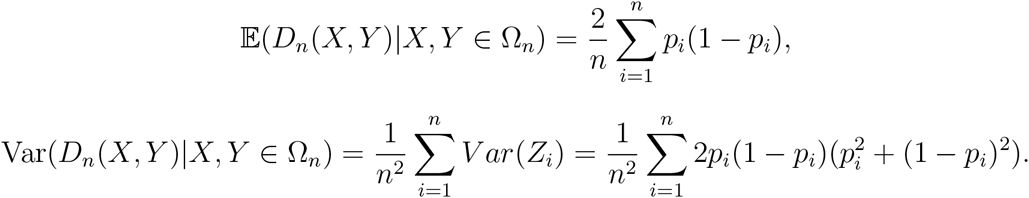

**Corollary B.3.1.** *The variance of the normalized Hamming distance between two genotypes will approach to zero with rate* 1/4 *n as n →* ∞, *i.e. there is an equidistance property as n large for the set of total sampled genotypes.*

*Proof*. The statement follows from

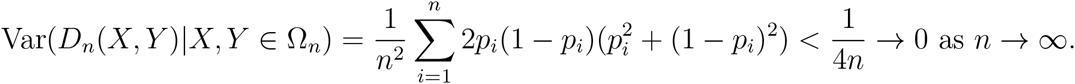

This explains that when *n* large enough, even though the portion of the typical genotypes is small, the normalized Hamming distance between two genotypes is close to the normalized Hamming distance of two (*n,ε*)–typical genotypes.

Now, given *ε* > 0 and 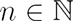, we denote by 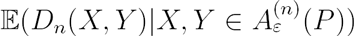 the average normalized Hamming distance of two typical genotypes. Then

**Proposition B.3.2.** *The following estimates holds for n large enough,*

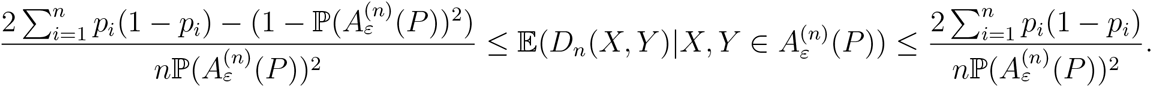

*Proof*. We note that for *n* large then 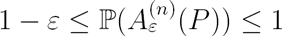. Therefore

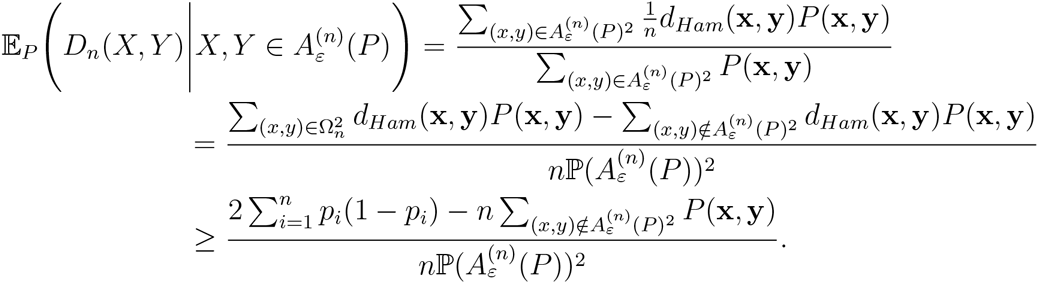

It implies the proof.

We then immediately have following corollaries:

**Corollary B.3.2.** *We have for n large*

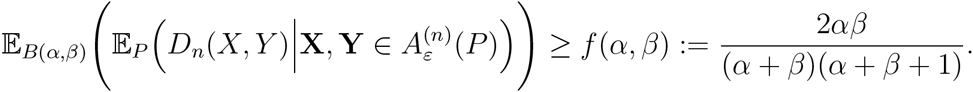

**Corollary B.3.3.** *This lower bound f*(*α, β*) *is monotone along the average entropy rate 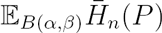. It means that when the average entropy rate increases then the below bound f*(*α, β*) *increases and vice verse.*

We also have a nice following property

**Theorem B.3.1.** *The correlation between the absolutely difference of logarithm with base 2 of probabilities of two arbitrary genotypes and their Hamming distance is always non-negative, i.e.*

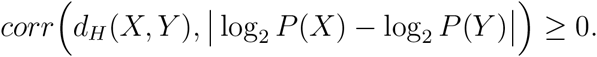

*Proof*. First, by denoting

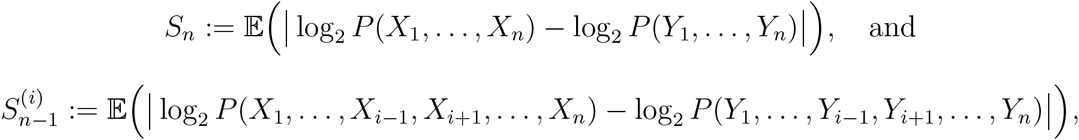

it is easy to see that

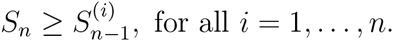

Indeed, we have (for shorting the notations, we use here 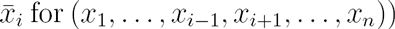

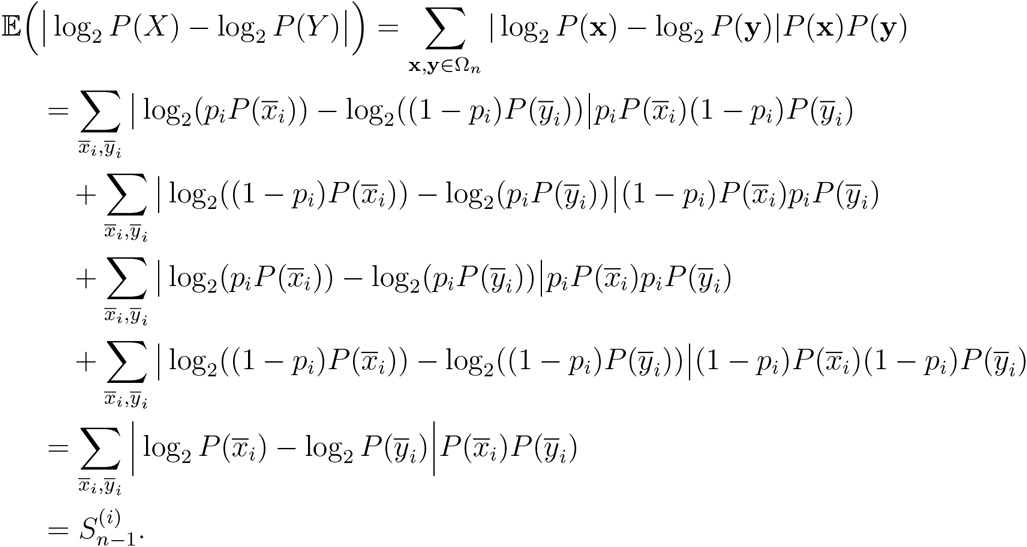

Therefore,

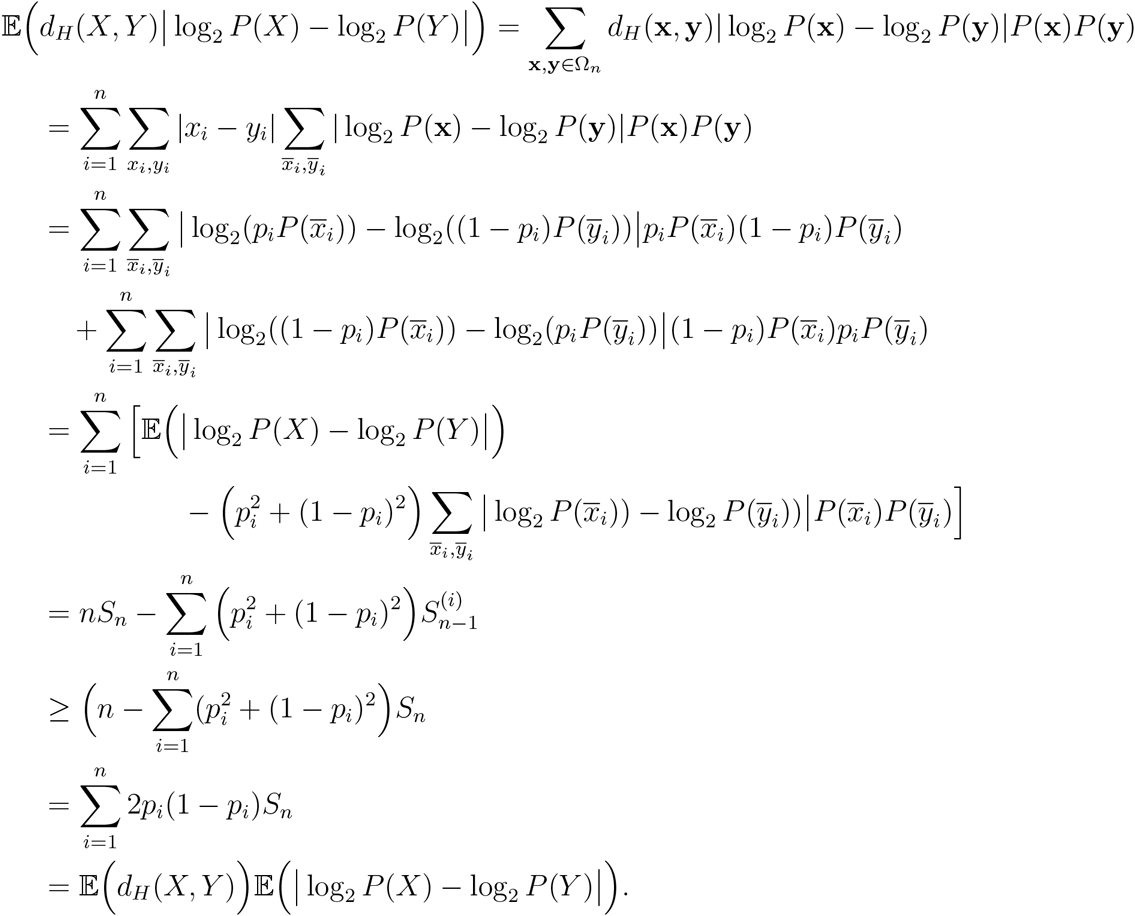

This implies the proof.

### B.4 Non-stationary AEP

In this section of the Appendix, we consider some AEP properties in the non-stationary case:

**Proposition B.4.1.**

1. *Given a sequence of binary independent random variables* {*X*_*n*_} *with the corresponding mass probability functions p*_*n*_(·) *sastifying*

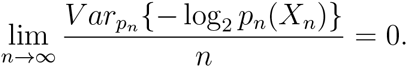 *Then, we have*

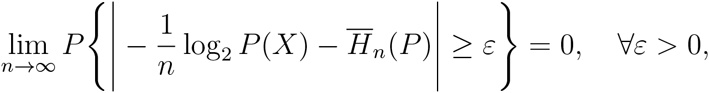

*where 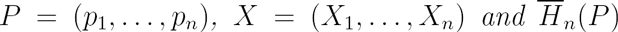 is the entropy rate with respect to P*.
2. *Given a sequence of binary independent random variables* {*X*_*n*_} *with the corresponding mass probability functions 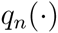 sastifying 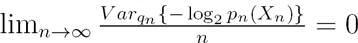. Then, we have*

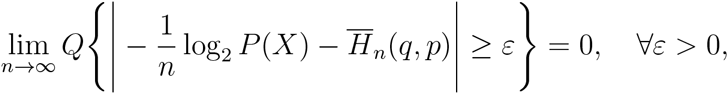

*where 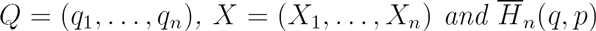 is the cross entropy rate of Q with respect to P.*

*Proof*. We will prove the second statement. The first one can be done similarly. Indeed, we have

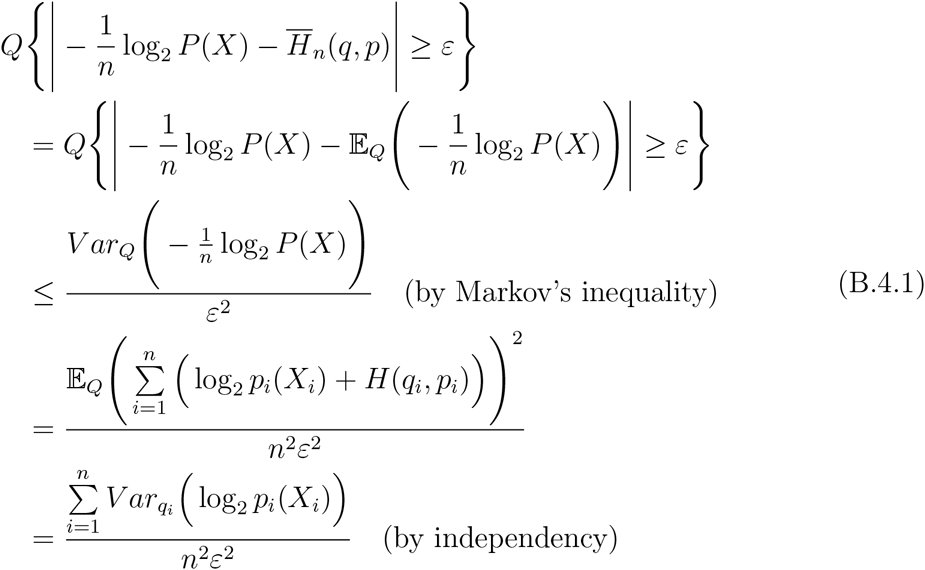

Therefore we obtain

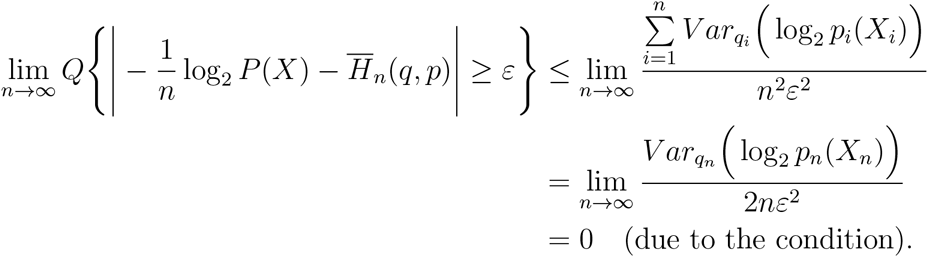

It implies the proof.

**Proposition B.4.2.** *Let* {*X*_*n*_}_*n*_ *be a sequence of mutual independent random variables with given binomial distribution 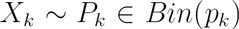. Given any other sequence of binomial distributions 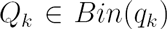 with assumption that 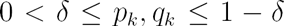 for all k. Then*

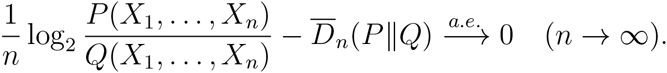

*Proof*. Denote by 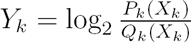 and its sample average 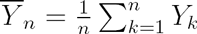. Note that

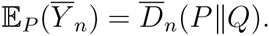

Moreover, from the assumption of *p*_*k*_, *q*_*k*_ we have

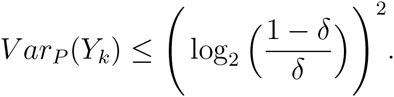

Therefore by applying the strong law of large numbers we obtain the result.

## C Appendix C

### C.1 Quantitative versions of the AEP

In this section of the appendix, we will show the following quantitative versions of the AEP and the cross entropy AEP. For all *ɛ* > 0 and 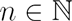, it holds that

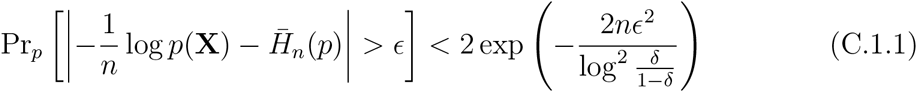

and

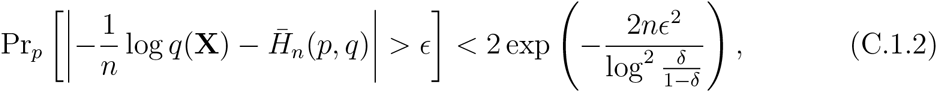

where by Pr_*p*_ we denote the probability given that the genotype X is distributed according to *P*.

These estimates can be obtained as follows. Suppose *Z*_1_,…, *Z*_*n*_ are independent, realvalued random variables, with *Z*_*i*_ taking values in the interval [*α*_*i*_, *b*_*i*_]. Then the Hoeffding inequality states that

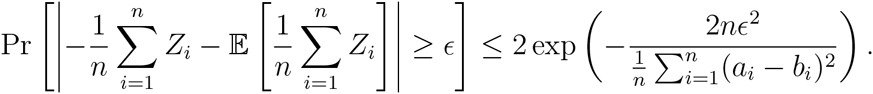

First, we apply the Hoeffding inequality to the random variables *Z*_*i*_ taking on the value − log *p*_*i*_ with probability *p*_*i*_, and the value − log(1 − *p*_*i*_) with probability (1 − *p*_*i*_). The Hoeffding inequality then implies

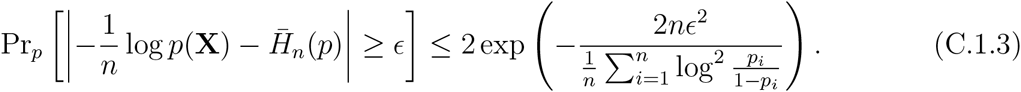

Similarly, we could define *Z*_*i*_ to be equal to −log *q*_*i*_ with probability *p*_*i*_ and equal to − log(1 − *q*_*i*_) with probability (1 – *p*_*i*_). Then, the Hoeffding inequality reads

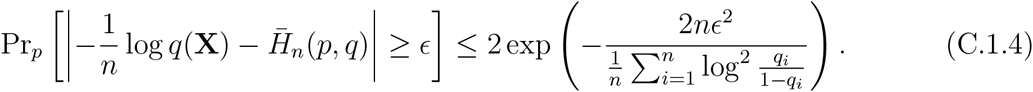

Note that the above inequalities can be viewed as versions of the AEP with explicit, exponential error bounds, for non-stationary sources.

### C.2 Error bounds for typicality classifiers

In this section we explain how the quantitative versions of the AEP from the last section imply exponential error bounds for the typicality classifiers introduced in the main text.

#### C.2.1 Error bound for naive typicality classifier

We assume without loss of generality that 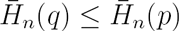. We recall the definition of the constants

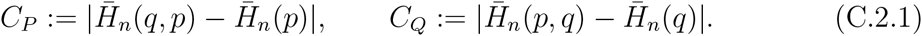

and the definition of the error rate

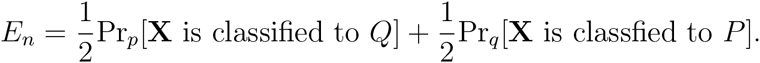

We note that in the naive typicality classifier, given that a sample **X** comes from *Q*, an error can only be made, that is it can only be assigned to *P*, if

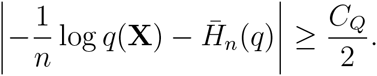

The quantitative AEP bounds the probability of this event by

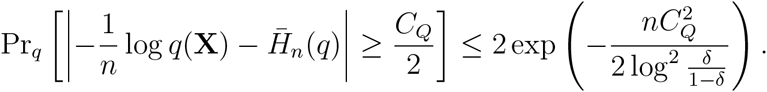

Given that a sample is drawn from *P*, an error can be made in two situations, either

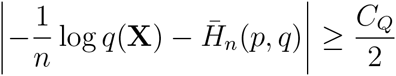

or

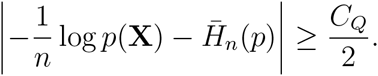

The quantitative cross entropy AEP bounds

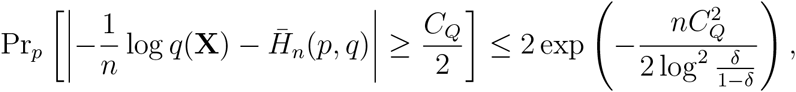

whereas the quantitative AEP implies

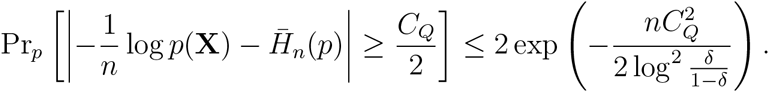

Consequently,

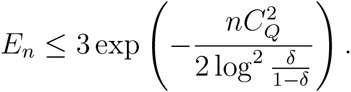

#### C.2.2 Error bound for cross entropy classifier

We now assume without loss of generality that *C*_*Q*_ > *C*_*P*_. Note that given that a sample **X** comes from distribution *Q*, it can only be assigned to *P* if

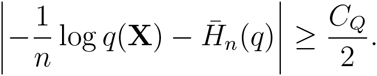

As in the previous section, the quantitative AEP bounds this probability of this event by

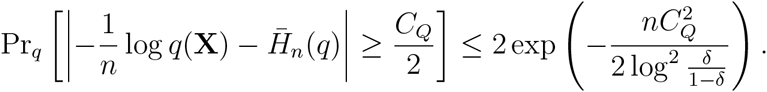

Similarly, given that a sample **X** comes from distribution *P*, it can only be assigned to *Q* if

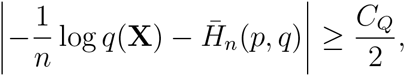

and the quantitative cross entropy AEP estimates

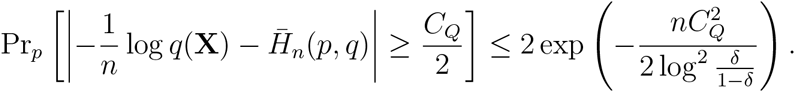

Combining these two estimates we obtain

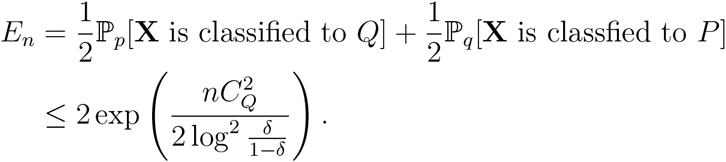

In fact, by using one-sided Hoeffding inequalities (and corresponding one-sided AEPs), one can actually replace the prefactor 2 by 1.

### C.3 Domain in log-probability plane

In this section we consider the limiting behavior for *n →* ∞ of the sets 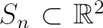 which we define by

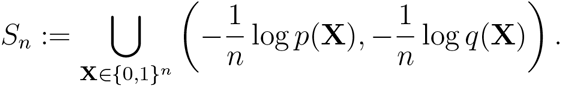

These sets are the union of the image of all possible genotypes in the log-probability plane.

The claim is that (with probability one) these sets converge (in Hausdorff distance) to a certain closed, convex set *A*. This set *A* is determined by the distribution of the *p*_*i*_’s and the *q*_*i*_’s. Loosely speaking, for large *n*, for every point *A* there is a point in *S*_*n*_ closeby, and for every point in *S*_*n*_ there is a point in *A* closeby.

For simplicity, we assume that the gene frequences *p*_*i*_ and *q*_*i*_ can only attain a finite number of values. We denote the possible values for *p*_*i*_ by a_1_…, *a*_*N*_ and the possible values for *q*_*i*_ by *b*_1_,…,*b*_*N*_. We assume moreover that 0 < *a*_1_ <…< *a*_*N*_ < 1 and 0 < *b*_1_ <…< *b*_*N*_ < 1.

We denote by *f*(*α*_*j*_, *b*_*k*_) the probability that *p*_*i*_ = *a*_*j*_ and *q*_*i*_ = *b*_*k*_.

By *L*(*a, b*) we denote the (unoriented) line segment between the points (−log(*a*), −log(*b*)) and (−log(1 − *a*), − log(1 − *b*)). Then the set *A* is the Minkowski linear combination of the line segments *L*(*a*_*j*_, *b*_*k*_), that is

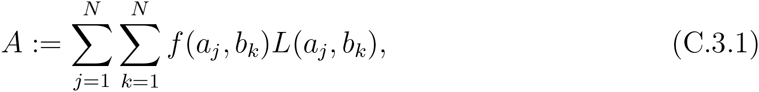

where the sums on the right-hand-side denote Minkowski sums.

**Theorem C.3.1.** *With probability 1, the sequence of p_i_ and q_i_ is such that the set S_n_ converges to the set A in the Hausdorff distance as n →* ∞.

A version of this theorem is also true when *p*_*i*_ and *q*_*i*_ are continuously distributed, under some extra conditions on the distribution (specifically their behavior close to 0 and 1). The set *A* then has a description as a ‘Minkowski integral’ rather than a Minkowski sum. We do not focus on this case to avoid technicalities.

The Hausdorff distance between two bounded and closed sets *K*_1_ and *K*_2_ is defined as the smallest *ϵ* ≥ 0 such that *K*_1_ is contained in *T*_*ϵ*_(*K*_2_) and *K*_2_ is contained in *T*_*ϵ*_(*K*1), where

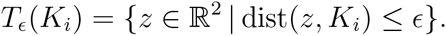

We will explain the proof of the theorem. We let *N*_*n*_(*a*_*j*_, *b*_*k*_) denote the number of indices *i* ∈ {1,…, *n*} such that *p*_*i*_ = *a*_*j*_ and *q*_*i*_ = *b*_*k*_.

For the first part of the proof, we define auxiliary sets *A*_*n*_ by

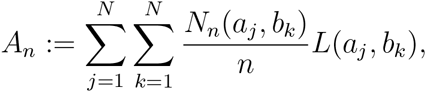

and we will show that *A*_*n*_ *→ A* in the Hausdorff distance. For instance by Sanov’s theorem, it follows directly that with probability 1,

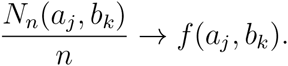

By the continuity properties for the Minkowski sum it follows that the sets *A*_*n*_ converge in the Hausdorff distance to *A.*

With a bit more work (and an application of for instance Pinsker’s inequality and the Borel-Cantelli Lemma), one can also extract that with probability one, the convergence is faster than 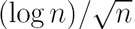.

In the second part of the proof, we show that the Hausdorff distance between *A*_*n*_ and *S*_*n*_ can be bounded by *C*/*n*, for some constant *C*. In fact, we will see that *A*_*n*_ is the convex hull of *S*_*n*_, while on the other hand *S*_*n*_ is a *C/n-net* in *A*_*n*_, which means that for every point in *A*_*n*_, there is a point in *S*_*n*_ at distance less than *C/n*. First, we introduce some additional notation.

For a line segment *L* in 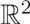, we denote by *B*(*L*) and *E*(*L*) its endpoints, in such a way that *B*(*L*)_2_ ≤ *E*(*L*)_2_, and if *B*(*L*)_2_ = *E*(*L*)_2_, then *B*(*L*)_1_ ≤ E(*L*)_1_. These conditions uniquely define *B*(*L*) and *E*(*L*).

We will now give an equivalent description of the set *S*_*n*_. We start with an important observation. Given a string **X** ∈ {0, 1}^n^, the point

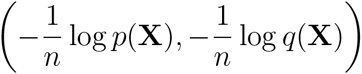

only depends on for how many indices *i, X*_*i*_ = 1 and *p*_*i*_ = *a*_*j*_, *q*_*i*_ = *b*_*k*_. This motivates the following definition.

By *M*^*n*^ we denote the space of *N × N* matrices *x* with integer entries that satisfy the constraints

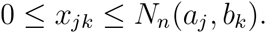

For *x* ∈ *M*^*n*^ we denote by 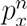 the following point in 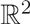

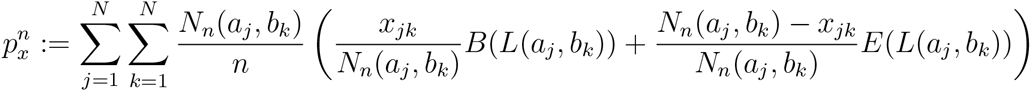

It is then clear that we may rewrite *S*_*n*_ as

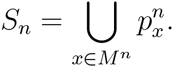

Moreover, it follows that *S*_*n*_ ⊂ *A*_*n*_.

Using this representation of *S*_*n*_, we can now check that as *n →* ∞, the Hausdorff distance between *S*_*n*_ and *A*_*n*_ is bounded by *C/n*, thereby proving the theorem.

A line segment is the convex hull of its endpoints. For two sets *B*_1_ and *B*_2_, the convex hull of *B*_1_ + *B*_2_ is equal to the convex hull of *B*_1_ plus the convex hull of *B*_2_. Therefore, the set *A*_*n*_ is equal to the convex hull of the Minkowski sum

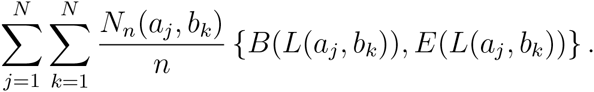

In other words, if we denote by *M*_*N*_ the set of all N × N matrices with entries either zero or one, the set *A* can also be described as the convex hull of the points

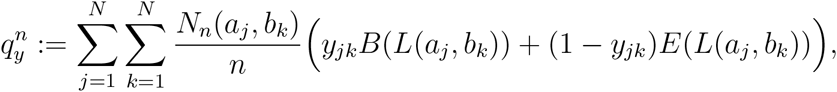

for 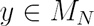, that is

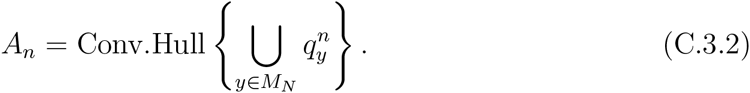

Note that the set 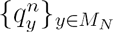 is a subset of 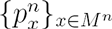, while we established previously that 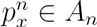 for every 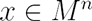. Hence, also

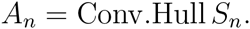

The final statement to check is that every point in *A*_*n*_ is within distance *C/n* to some point 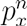. Let therefore *α* ∈ *A*_*n*_. Then

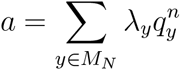

for some constants *λ*_*y*_ ≥ 0 with 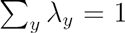. If we plug in the definition of 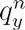 and switch the order of summation, we may write *α* as

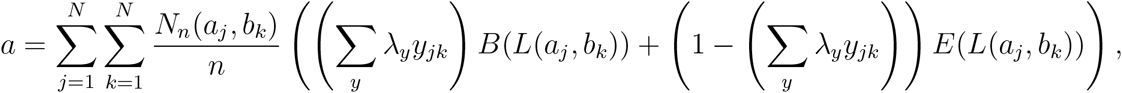

where we used that *∑*_*y*_ *λ*_*y*_ = 1. Then choose *x*_*jk*_ such that

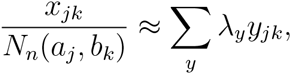

the error being bounded by at most 1/*N*_*n*_(*α*_*j*_, *b*_*k*_).

The distance between *a* and

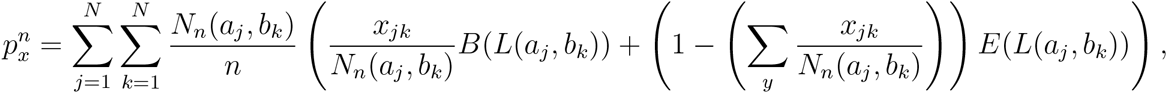

is therefore bounded by *C/n* for some constant *C* depending on *N* and the distance of the *a*_*j*_ and *b*_*k*_ to 0 and 1. This finishes the proof of the theorem.

#### C.3.1 A practical method to compute the accessible set *A*

The previous description (C.3.2) provides a way to compute the set *A*_*n*_ and a similar formula can be derived for *A*. However, it is not very efficient. In this section we will provide a more efficient way to calculate *A*, by specifying its boundary.

First we order the points (*a*_*j*_, *b*_*k*_) according to the angles

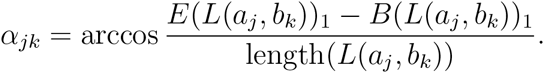

In other words, for *ℓ* = 1,…, *N*^2^, we let *j*(*ℓ*) and *k*(*ℓ*) be such that

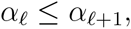

where we used shorthand 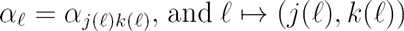 is surjective onto {1,…, *N*}^2^.

Next, with obvious abbreviations, we define vectors

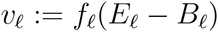

and

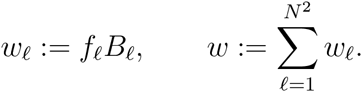

It is immediate from the definitions that the set *A* can also be written as

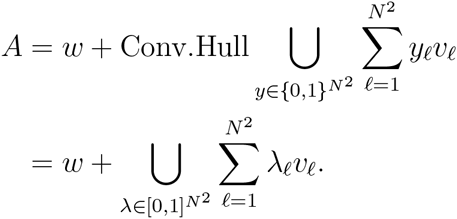

We claim that

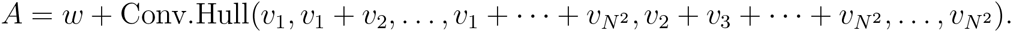

To see this, we first note that we may without loss of generality assume that *w* = 0, and that the slopes of 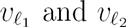 are different when 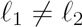.

By the definition of *B*_*ℓ*_ and *E*_*ℓ*_, we know that for every *ℓ*, the vector *υ*_*ℓ*_ either points to the right or lies in the upper halfplane. Note that the origin lies in *A*, as do the line segments 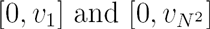. Moreover, the set *A* lies in the smaller cone bounded by the rays starting from the origin with the directions of 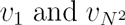 respectively. It follows that the origin is an extreme point of the convex polyhedron *A.*

Note that for *k* = 1,…, *N*^2^ – 1 we may alternatively write *A* as

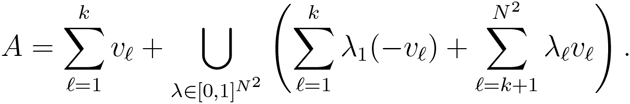

This representation of *A* allows one to check that for every *k =* 1,…, *N*^2^,

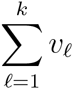

is an extreme point of *A*, while the line segments

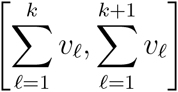

are faces of *A*. Indeed, it is clear that the points and line segments lie in *A*. On the other hand, *A* is contained in the smaller cone bounded by the rays with starting point

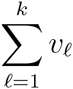

and directions 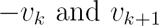 respectively. A similar argument shows that the points

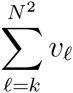

are extreme points and the line segments

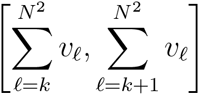

are faces. Hence, we have shown that

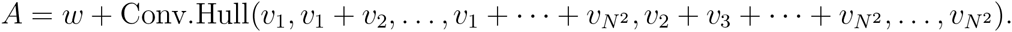

This description allows for fast checks whether or not a point lies in *A*.

## Footnotes

1 Note that in probability theory, the entropy rate or *source information rate* of a stochastic process is defined *asymptotically*, 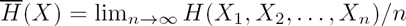 when this limit exists.

2 We may also explicitly express the error rate of this classifier in a closed form (Appendix A.1).

3 As with the naïve typicality classifier, we may explicitly express the error rate of this classifier in a closed form (Appendix A.2).

4 Under a particular *restrictive assumption* on the underlying SNP frequency model and for large enough n, the classifier may use the entropy rates as proxy, due to the following *asymptotic* result, 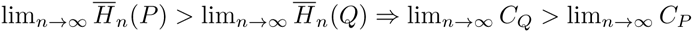 (Appendix B, Corollary B.2.2)

5 A natural estimator, which simply counts the proportion of alleles of a particular type, and a maximum likelihood estimator (MLE) give identical solutions when the sample consists of unrelated individuals. Thus maximum likelihood provides a justification for using the “natural” estimator ([Adrianto and Montgomery, 2012]).

6 The performance of the typicality classifiers under MLE can also be formally captured (Appendix A.3).

7 The standard approach is to take the mean of the posterior distribution. The beta distribution is a conjugate prior for the binomial likelihood (which is our sampling distribution) since the posterior is also a beta distribution, making the formulation of the posterior simple: *Beta*(*z* + *α, N* – *z* + *β*), where *Beta*(*α, β*) is the prior, *N* is the size of the sample and *z* is the number of ‘1’ alleles in the sample at that locus [Schervish, 1995]. We then take the mean of the posterior which is (*z* + *α*)/(*N* + *α* + *β*).

8 Otherwise, we would have to incorporate the full information from the *joint* distribution of SNPs across loci, which is over and above the low-dimensional standard LD statistics.

9 In simulating *E*_*n,m*_ we replace allele frequency estimates of zero with a small constant, 1/(*m* + 1), a common procedure to avoid zero genotype frequencies ([Rosenberg, 2005]l [Phillips et al., 2007]).

